# “Prefrontal cortical contributions to working memory loading, maintenance and recall are parsed by hippocampal-prefrontal oscillatory assembly dynamics”

**DOI:** 10.1101/2021.12.20.473436

**Authors:** Aleksander P.F. Domanski, Michal T. Kucewicz, Eleonora Russo, Mark D. Tricklebank, Emma S.J. Robinson, Daniel Durstewitz, Matt W. Jones

**Affiliations:** School of Physiology, Pharmacology & Neuroscience, Faculty of Life Sciences, University of Bristol, University Walk, Bristol BS8 1TD, UK; The Alan Turing Institute, 96 Euston Road, London, UK; The Francis Crick Institute, 1 Midland Road, London, UK; BioTechMed Center, Multimedia Systems Department, Faculty of Electronics, Telecommunications and Informatics, Gdansk University of Technology, Gdansk, Poland; Dept. of Theoretical Neuroscience, Central Institute of Mental Health, Medical Faculty Mannheim, Heidelberg University, 68159 Mannheim, Germany; Johannes Gutenberg University, Mainz, Germany; Centre for Neuroimaging Science, King’s College London, Denmark Hill, London, UK

**Keywords:** systems neuroscience, working memory, prefrontal cortex, hippocampus, oscillations, population coding, decision making, DNMTS

## Abstract

Working memory enables incorporation of recent experience into subsequent decision-making. This processing recruits both prefrontal cortex and hippocampus, where neurons encode task cues, rules and outcomes. However, precisely which information is carried when, and by which neurons, remains unclear. Using population decoding of activity in rat medial prefrontal cortex (mPFC) and dorsal hippocampal CA1, we confirm that mPFC populations lead in maintaining sample information across delays of an operant nonmatch to sample task, despite individual neurons firing only transiently. During sample encoding, distinct mPFC subpopulations joined distributed CA1-mPFC cell assemblies hallmarked by 4-5Hz rhythmic modulation; CA1-mPFC assemblies re-emerged during choice episodes, but were not 4-5Hz modulated. Delay-dependent errors arose when attenuated rhythmic assembly activity heralded collapse of sustained mPFC encoding; pharmacological disruption of CA1-mPFC assembly rhythmicity impaired task performance. Our results map component processes of memory-guided decisions onto heterogeneous CA1-mPFC subpopulations and the dynamics of physiologically distinct, distributed cell assemblies.

## Introduction

Working memory’s capacity to integrate prior experience into context-dependent choice is a cornerstone of adaptive decision-making^1^. Delayed non-match to sample (DNMTS) paradigms hinge on such memory-guided decisions and are subserved by interactions between executive and mnemonic hub regions including the prefrontal cortex (PFC) and hippocampus^2,3^. PFC principal neuron spike rates during delayed response tasks encode diverse features of sample identity and task rules in both non-human primates^4–8^ and rodents^9–15^, with sustained PFC principal neuron firing offering an intuitive neural correlate of working memory maintenance during task delay phases^16,17,26–32,18–25^. However, as analyses have extended from individual neurons to simultaneously recorded cortical populations, other informative features of PFC ensemble dynamics have emerged^33–35^.

Sequentially active, task-informative neurons in PFC and interrelated regions can span the behavioral progression from sample to choice^24,36–39^, and recent models invoke dynamic changes to the information coded by individual neurons and subpopulations across sample, delay and response epochs^20,37,48,40–47^. Thus subsets of transiently active neurons not classically selective for individual task features can contribute to DNMTS performance^35,49–53^. Here, we test whether and how these complex neural dynamics within the PFC – which have been characterized primarily in primates – reflect its interactions with hippocampus, using simultaneous hippocampal-PFC recordings in rats.

Hippocampal CA1/CA3 single unit activity during DNMTS-related tasks in both macaque^54^ and rat^11,55^ show dissociable sample, delay and choice correlates, potentially reflecting interactions with PFC. Indeed, simultaneous recordings from rodent PFC and hippocampus reveal co-varying network activity associated with 5-10Hz ‘theta’ frequency coherence across the two regions during route choice on mazes^56–59^ and spatially-guided object memory retrieval^60^. However, the precise timing and nature of the network physiology driving these interactions remains equivocal, as do the relationships between single neuron and population-level contributions to information encoding and retrieval across the hippocampal-PFC axis. For example, how are hippocampal representations integrated into PFC dynamics during both cue sampling and subsequent choice? Do different oscillatory interactions with hippocampus disambiguate the cognitive contexts of sample and choice?

We examined the dynamic contributions of hippocampal and PFC populations to information encoding during the DNMTS task. Specifically, we tested the hypotheses that: (1) Correlated groups of neurons distributed across hippocampus and PFC form assemblies that contribute to the representation of cue information during sample encoding and recall; (2) Dissociable subsets of PFC neurons (less directly modulated by hippocampus) maintain cue information during the working memory delay; and (3) At least one of these population signatures should fail to encode, maintain or transfer information during errors, culminating in an incorrect choice.

## Results

### Dissociable dCA1 and mPFC population dynamics reflect differential contributions to information encoding, maintenance, and recall during the DNMTS task

Six rats were trained on the DNMTS task (**Fig. 1A** and **S1A**) and chronically implanted with tetrodes (**Fig. 1B**, **Fig. S1C-F**) to record simultaneous spiking activity from dorsal CA1 and prelimbic medial PFC (mPFC); data are presented from the final two DNMTS sessions after criterion had been achieved (**Methods**). 31±5 mPFC and 30±5 dCA1 (mean±SEM) well-isolated putative principal neurons with mean firing rates >0.5Hz were analyzed from each animal per session.

**Figure 1:**
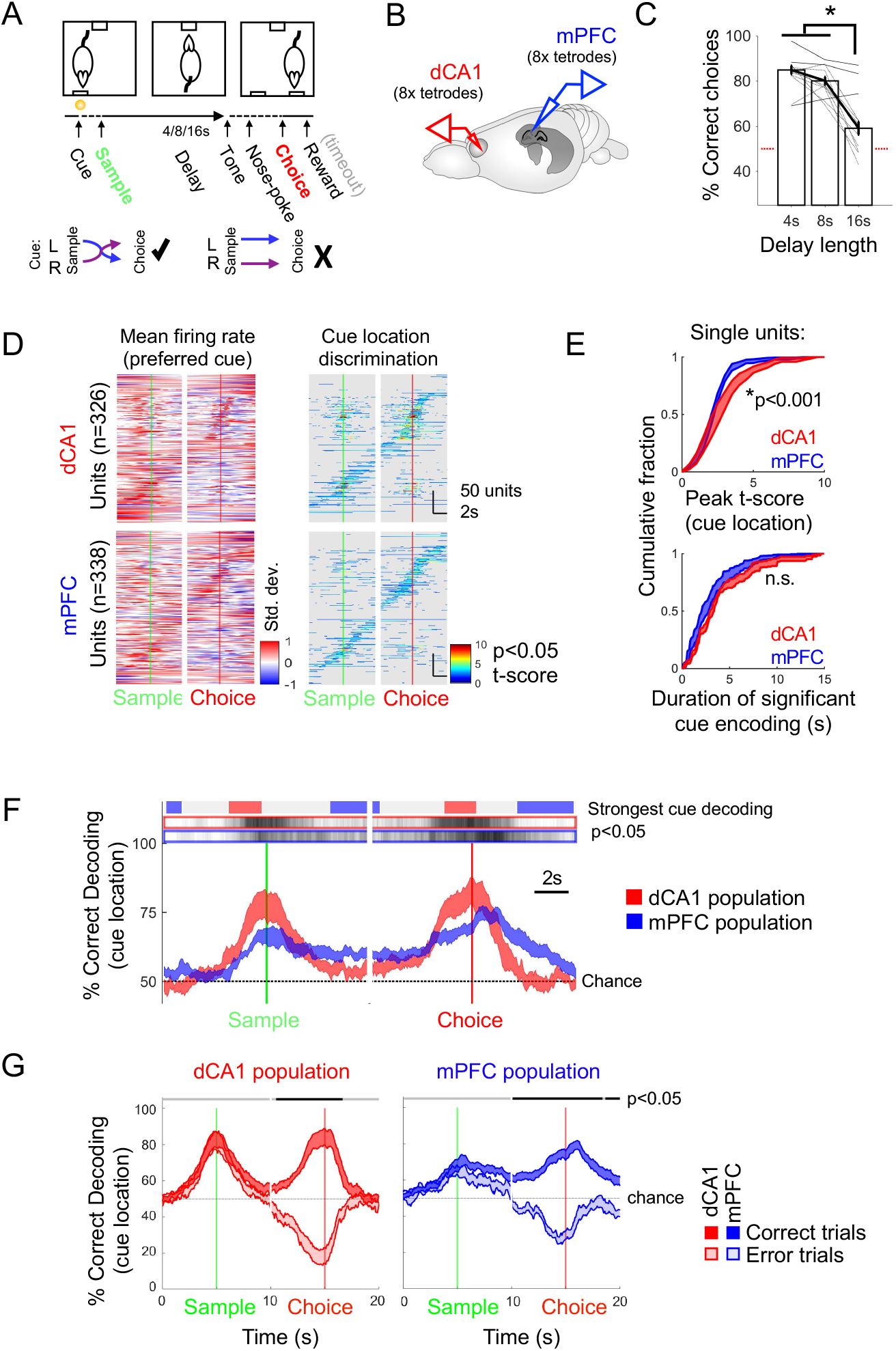
Differential contributions of dCA1 and mPFC neurons and populations to performance in the DNMTS working memory task **A** DNMTS task Schematic. Solid/dashed lines indicate periods of instructed/free-paced behavior. Incorrect choices led to a time-out before the subsequent trial. Below: Contingencies leading to Correct (left) and Error outcomes. **B** Schematic of chronic recording configuration. **C** Performance of rats on 4s, 8s and 16s delay trials. Black dotted/solid lines link individual trials from at/above chance sessions, red dotted line indicates chance level. **D** Left: Z-scored mean firing single units rates ±5s relative to Sample and Choice lever presses for preferred (higher peak firing rate) cue locations. Units from each brain area are sorted by times of peak discrimination of cue location. Right: time-resolved t-scores between cue location-specific firing rates. Grey regions mask periods of insignificant cue location discrimination (p<0.05, vs. Bootstrapped 95% CIs). Single units shown aggregated from all sessions. **E** Distributions of peak strength (top) and duration (bottom) of cue location encoding by dCA1 and mPFC single units. Shaded regions indicate mean±SEM of distributions across sessions. **F** Leave-one-out decoding of cue location from firing rates of single units in 50ms bins. Shaded curves indicate mean±SEM decoding across animals with matched trial and unit counts randomly sampled from available data. Grey shaded bars above indicate times of cue decoding periods significantly different from chance (cue-shuffled data). Blue/Red bars show times of significantly stronger cue decoding from mPFC/dCA1 units (p<0.05, bootstrap permutation test between the two conditions n=12 sessions). **G** Cross-validated decoding of cue location from dCA1 and mPFC units on error trials: Performance of regularised linear decoder trained on correct trials and tested on correct (black) and error (red) trials. Mean±SEM performance across recording sessions shown. Black bars indicate times of significant drops in cross-validated decoding performance (Bootstrap permutation test, Bonferroni-corrected p<0.05).

Every trial began with a sample phase initiated by light-cued presentation of either the right or left lever. Following sample press, the lever was retracted and rats turned and waited at a food pellet receptacle in the opposite wall; a tone then signalled the end of a 4, 8 or 16s delay, varying randomly from trial to trial to discourage mediating behavior. The choice phase was initiated by nose-poking inside the receptacle to release presentation of the two levers on the opposite wall. Correct choice was rewarded according to a nonmatch rule.

Rats made significantly more errors on 16s delay trials than trials with shorter delays (**Fig. 1C**, N=12, 2 sessions from 6 rats, ANOVA, F(2,36) = 42.4, p<0.001, Tukey-Kramer post-hoc test for delays p<0.001): all rats performed the DNMTS task significantly better than chance on 4s and 8s delay trials in both of two consecutive test sessions (p<0.05, binomial tests for each rat’s performance), whereas only three rats achieved above-chance performance at 16s delay in the latter of the two sessions. Response latencies did not differ systematically across 4, 8 or 16s delay trials, or correct vs. error trials (**Fig. S1B**), indicating that inaccurate performance on 16s delay trials was unlikely to stem from failure to engage with the task.

To quantify the time-varying encoding of cue location by single units, we used Student’s *t*-statistic as a measure of discrimination between left-*vs*. right-cue firing rates, tracking in 50ms bins how much information about the sample lever identity was carried by each neuron’s trial-averaged activity in dCA1 (**Fig. 1D**, top) and mPFC (**Fig. 1D**, bottom). For dCA1 units, discrimination tended to peak around sample and choice lever presentations. The activity of mPFC units was less bound to the sample and choice lever presses, and tiled the entire delay period in sequential, overlapping fashion (**Fig. 1D**, bottom right panel and **Fig. S1H-J**). Indeed, whilst dCA1 units showed significantly stronger peak cue location encoding than mPFC units during the sample and choice-preparatory periods (**Fig. 1E**, top, peak t-score, dCA1 vs. mPFC units; 3.49±0.23 vs. 2.72±0.08, t(463)=3.52, p=0.00048, t-test, N= 223, 242 units from 12 sessions), single units from the two areas were indistinguishable in the durations over which they encoded cue location more than expected by chance (**Fig. 1E**, bottom, duration of encoding, dCA1 vs. mPFC units; 2.24±0.19s vs. 1.99±0.16s; t(463) = 1.38, p=0.17, t-test, N= 223, 242 units from 12 sessions). Very few mPFC units showed persistent lever-selective delay firing (approximately 85% units showed significant decoding for <6s, **Fig. 1E**, bottom).

We next quantified how joint activity of simultaneously recorded populations of single units within each area carried task-relevant information using a linear discriminant classifier (**Fig. 1F**). Independently for each session, we used leave one out cross-validation (LOOCV) to predict lever identity for each 50ms time bin. To compare between animals, random subsets of equalized unit numbers and trial counts were drawn between conditions, ruling out dimensionality confounds in classifier performance.

Joint activity of simultaneously recorded dCA1 units showed strong but transient read-out of sample identity, peaking around the lever-presses but dropping to chance decoding performance during the delay period (**Fig. 1F**, red traces). Meanwhile, the discrimination around sample and choice lever presentation was less pronounced in mPFC populations, which instead maintained a stronger representation of sample identity than dCA1 populations throughout the delay and post-choice evaluation period (**Fig. 1F**, blue traces). These findings are in good agreement with recent comparisons between task coding dynamics in hippocampal and frontal cortical populations in primates^61,62^.

Which features of the dCA1 and mPFC population activity are essential to the correct execution of the DNMTS task? Previous studies have approached this question by inducing forced errors with lesioning or inactivation of the hippocampal-frontal network (O’Neill et al., 2013; Spellman et al., 2015), but less is known about the system’s dynamics during spontaneous, unforced errors. Since 16s delay trials challenged the working memory limits of rats, we quantified which aspects of the sequential contributions of dCA1 and mPFC populations failed during incorrect choices.

Decoders trained on correct trials and tested on error trials demonstrated that decoding of sample lever identity from hippocampal populations was intact (**Fig. 1G**): on error trials the, two conditions were indistinguishable around the Sample lever press and dCA1 continued to represent the wrong (same) lever on approach to the Choice lever press. Thus, even as the rats re-visited the Sample lever, hippocampal representations remained faithful. In the mPFC however, sample representation on error trials began to decay immediately after the Sample lever press, such that incorrect choices could be predicted approximately 2s earlier in the delay period than from dCA1 activity (black bars in **Fig. 1G** indicate significantly error-predicting periods).

### mPFC population dynamics support coding that spans the DNMTS delay phase

What ensemble mechanisms might underlie the sustained coding of cue identity by mPFC populations during the delay period on correct trials? We examined data on a trial-by-trial basis to find associations between mPFC population dynamics and DNMTS accuracy (**Fig. 2**), comparing LOOCV decoding across delay lengths and correct *vs*. error outcomes. Subsets of the different trial types were drawn at random to allow matching of trial numbers across conditions (accounting for fewer available correct trials on longer delays).

**Figure 2:**
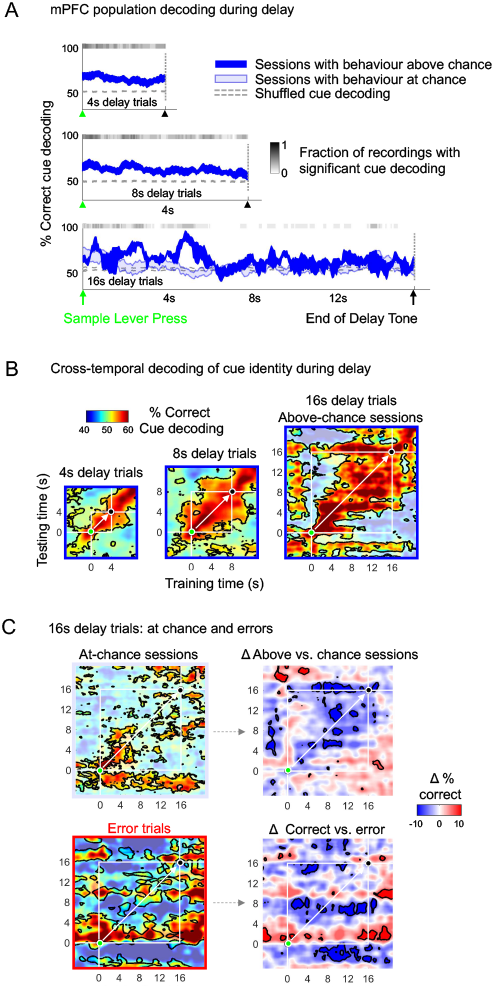
Maintenance of cue location by a stabilizing population code in mPFC underlies correct performance in the DNMTS task **A** Time-resolved decoding of cue location from mPFC single unit populations on correct trials for sessions above (solid blue) and below (light blue) chance performance (50%, dotted line). Fraction of above-chance performing sessions with decoding greater than 95% of cue location-shuffled bootstrap decoding distribution are shown as grey shaded bars above the curves. Mean±SEM decoding performance from 12 sessions (all above chance performance for 4,8s delays, nine at chance for 16s delay trials). **B** Cross-temporal decoding of cue location from populations of mPFC single units during the delay period: Cross-validated regularised linear decoders trained and tested at different points during the delay period (±5s) White lines indicate bounding box of delay period. Green and black markers indicate Sample press and end-of-delay tone, respectively. White arrow along diagonal indicates training and testing at the same time point (using different withheld trials for testing), providing curves shown in A. Bounded regions show significant (p<0.05) decoding relative to cue-shuffled bootstrap distribution. **C** Cross-temporal cue decoding performance of mPFC unit population recordings on correct 16s delay trials from at-chance sessions (top) and on errors on above-chance (bottom). Statistics as for B. Right: Subtraction of above-chance session decoding from below-chance sessions (top) and correct from error trials (bottom). Bounded regions indicate significant differences between conditions (bootstrap permutation test, p<0.05).

On 4s and 8s delay trials, during which all rats performed above chance in both recording sessions (**Fig. 1B**), delay period decoder results were significantly better than chance performance (bootstraps with shuffled trial labels) for the entire delay period duration in the majority of recording sessions (indicated by grey shading in **Fig. 2A**). For 16s delay trials, we split recording sessions by whether animals’ behavioural performance was at, or significantly better than chance performance, (light and solid blue curves in **Fig. 2A**). Here, whereas correct outcome 16s delay trials from above-chance sessions showed decoder performance that was above chance for the majority of the delay period (albeit variable due to the small subset of three above-chance sessions), mPFC population decoding from correct trials from chance-performance behavioural sessions fluctuated around chance levels from shortly after sample lever press. Thus, even though all trials examined corresponded to “correct” outcomes, faithful decoding of cue identity from mPFC populations demanded that rats were not guessing their choices, implicating faithful cue representation by mPFC populations in successful task performance.

Thus on correct trials, a delay-spanning coding scheme is implemented in mPFC during the working memory period of the DNMTS task, despite individual neurons firing only transiently (**Fig. 1D**); what form does this scheme take? One possibility is that firing rates across neurons evolve in fixed proportions relative to one another, such that a decoder trained on population firing rates at the start of the delay successfully predicts left *vs*. right sample lever identity using firing rates from the end of the delay, and vice versa. Alternatively, a dynamic code implemented by the mPFC population may mean decoding results are only transiently valid around the time of the training data. We compared evidence for these two hypothetical schemes by constructing decoders using population firing rate vectors from each 50ms segment of the delay period and systematically ‘sliding’ the test data across the entire delay period (**Fig. 2B**, method reviewed in Meyers et al., 2008). These results form cross-temporal decoders on symmetrical axes, on which the diagonal (white arrows in **Fig. 2B**) represents training and testing performed at matching time-points. We trained and evaluated separate decoders for each recording session; results in **Fig. 2B** summarise mean performance across sessions.

For 4s and 8s delay trials, decoders trained and tested at any pair of times during the delay period were similarly effective, generating symmetrical regions of significant decoding extending throughout the delay (filling the white bounding boxes in **Fig. 2B**). However analysis of 16s delay trials from sessions with above-chance behaviour, revealed that mPFC populations encode the cue identity dynamically during the extended delay period. Significant off-diagonal decoding was observed during the first 8s of delay, but decoders trained on firing rates at the start of the delay were no longer successful when tested on firing rates from after 8s: whereas decoders trained at 8s or later successfully recovered cue identity until the end of the delay.

This sustained representation was seen neither in dCA1 (**Fig. S2**), nor in decoding results from sessions with at-chance behavioural performance (**Fig. 2C**). Instead, despite an initial early-delay period of transient decoding comparable in strength to the above-chance sessions (as in **Fig. 2B**), cross-temporal decoding did not outlast approximately 2s. These results link cue encoding in mPFC populations to successful DNMTS task performance. Consistent with this, the sustained mPFC population code evident during correct 16s trials was not established during error trials (**Fig. 2C**, lower panels).

### A subset of neurons with distinct rhythmic firing signatures form joint dCA1-mPFC cell assemblies recruited during sample and choice events of the DNMTS task

Thus far, our results demonstrate that dCA1 neurons are particularly informative during encoding (sample) and recall (choice), whereas mPFC populations are critically involved in delay phases, through the transient contributions of individual mPFC neurons to sustained population coding. Correct activation of the dCA1-mPFC pathway is essential for performance of spatial working memory tasks^64–66^, but given the dissociable encoding of task-related information in dCA1 and mPFC populations, when and how is information shared between the two regions during delayed responding?

We used factor analysis (FA)^67,68^ to detect coordinated activity among units from mPFC only, dCA1 only, or jointly from mPFC and dCA1 (**Fig. 3**, **Fig. S3**, **Methods**). In contrast to Principal Component Analysis (PCA) which detects variance-maximizing dimensions, FA is a model-based statistical tool that explicitly captures correlations between variables through a set of independent factors. FA has thus been shown to outperform PCA for the present purposes^69^. Recorded units were assigned to the same cell assembly when they significantly loaded on the same latent factor (**Fig. 3B-C**), which captured correlated firing rate activities within the population of neurons (**Fig. 3D**). Time-varying factor scores associated with each extracted factor in the FA model (**Fig. 3B**) measure assembly activation strength, as exemplified by significant event-locked activation of inter-area assemblies binding dCA1 and mPFC units during the DNMTS task (**Fig. 3D, 4A**, further examples in **Fig. S4A**).

**Figure 3:**
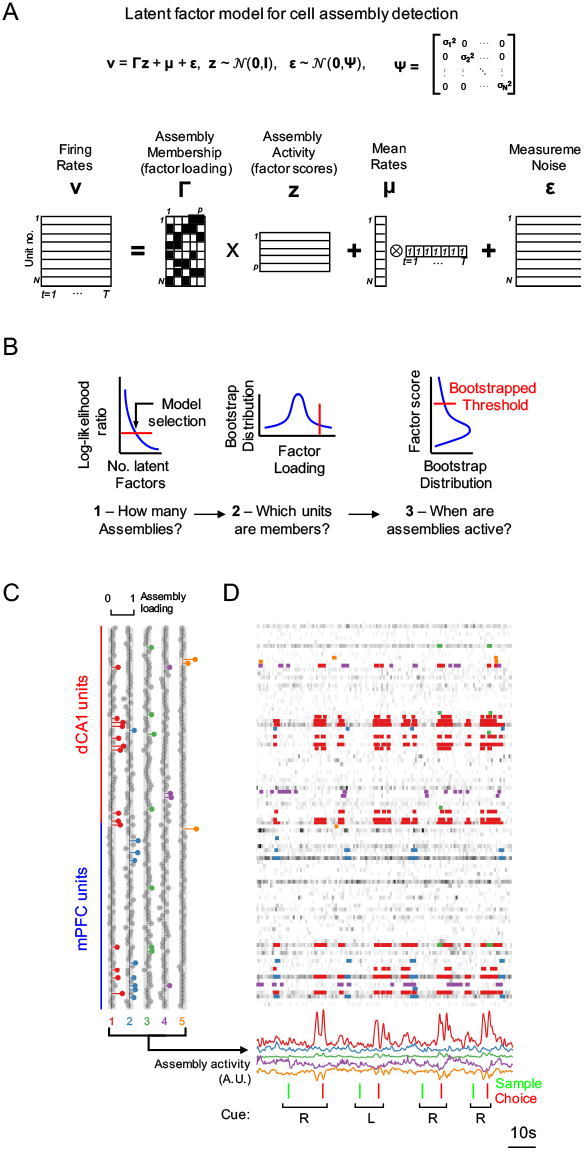
A latent factor analysis model detects correlated inter-regional dCA1-mPFC cell assemblies **A** Schematic illustration of the FA–based cell assembly detection method, decomposing parallel recordings from N single units into time-varying activation scores of p (p<N) factor scores. **B** Models selection steps involved in FA-based cell assembly detection. **C** Inter-area cell assembly detected from parallel dCA1-mPFC recording. Example loading of single units to the five detected latent factors (cell assemblies). Grey circles indicate units with insignificant loading strengths to the factors (shaded columns indicate p<0.01 vs. shuffled bootstrap distributions). **D** Example spike train raster from units in C. Below: Factor activation scores of the detected cell assemblies spanning dCA1 and mPFC (within-area assemblies not shown) during four successive trials in the DNMTS task. Colour annotation on spike trains indicates times of significant activation (bootstrap p<0.01, vs factor model calculated from shuffled spike rates) for each detected inter-area assembly.

**Figure 4:**
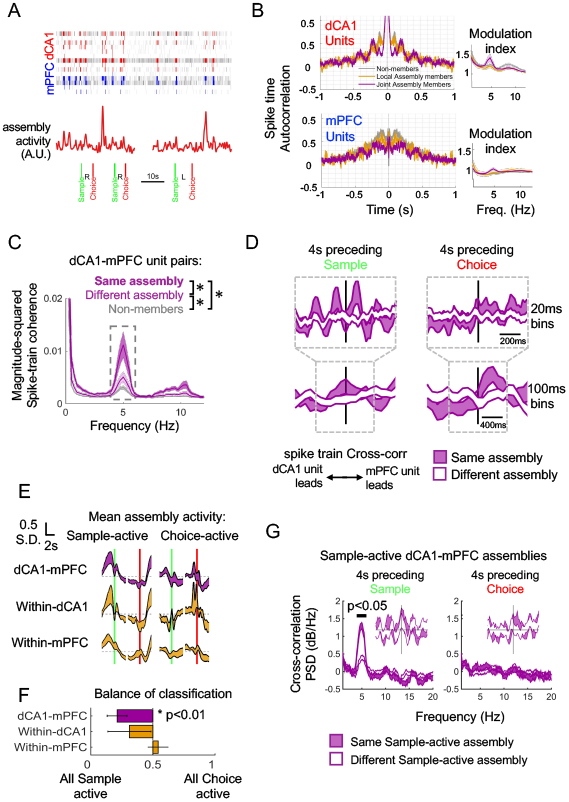
A subset of neurons with distinct rhythmic firing signatures form dCA1-mPFC-spanning cell assemblies which are active around the lever press events. **A** Spike trains of units comprising Assembly 1 (red annotation in Figure 3 C,D) highlighted to show firing of dCA1 and mPFC (red and blue, respectively) units during significant assembly activation times. **B** Left: Rate-normalized spike-time auto-correlation functions (mean±SEM across units) for nonassembly units, members of within-area and inter dCA1-mPFC cell assemblies for each area. Right: Modulation index of spike train autocorrelation functions. **C** Inter-area spike train coherence between dCA1-mPFC unit pairs within, across and outside cell assemblies (mean±SEM of dCA1-mPFC pairs across sessions shown). Grey shaded region indicates frequency range used for statistical comparison of rhythmic modulation (asterisks). **D** Pairwise dCA1-mPFC spike train cross-correlogram (top: 20ms bins, bottom: 100ms bins) for spikes fired by units in the 4s preceding Sample (left) and Choice (right) lever press events. Average across all pairs shown (mean±SEM across recording sessions) for within (purple) and across (white) assembly pairs. Random sampling of spikes (100 draws per cell pair) was used to match firing rate offsets between cells. **E** Standardized activation patterns (factor score activity) of different classes of cell assembly detected across all 12 recording sessions (mean±SEM across sessions), aligned to Sample and Choice lever press events. Assemblies were sorted by peak activity time and categories as either “Sample-active” or “Choice-active (Left and Right columns, respectively) based on strongest activation. Dashed lines are mean assembly activities for each class. **F** Fractions of cell assembly categorized as Sample- and Choice-active for each assembly type detected in the 12 recording sessions. 0 and 1 indicate assemblies were most strongly activated during the Sample and Choice phases respectively. Asterisk indicates that dCA1-mPFC assemblies were significantly biased towards more Sample-active. **G** Rhythmic 4-5Hz dCA1-mPFC spike correlations (as in D) were specific to pairs of units from Sample-active assemblies and were strongest for inter-regional pairs drawn from the same cell assembly.

Units from both dCA1 and mPFC that participated in cell assemblies spanning the two regions (**Fig. 3D**, **Fig. 4A**) showed 4-5Hz rhythmic modulation in the autocorrelation of their spike trains (**Fig. 4B**). dCA1-mPFC single unit pairs drawn from assemblies showed coherent spike train modulation at 4-5Hz, which was weaker for pairs drawn from different cell assemblies, and weaker again for pairs of units not detected as cell assembly members using FA (**Fig. 4C**: average 3.5-5.5Hz coherence across dCA1-mPFC cell pairs, F(2,27) = 50.0 p<0.0001, Tukey-Kramer post-hoc tests for assembly membership p<0.05, N = 2 sessions from each 6 animals, also see **Fig. S4B**). Coherent 4-5Hz spike train modulation was also weaker for pairs drawn from within-area cell assemblies in dCA1, and essentially absent between pairs from mPFC. This indicates that “non-assembly” units were physiologically distinct from their assembly counterparts. The 4-5Hz assembly motif was specific to the task period, since the spectral signature was absent from spike trains recorded during one hour rest periods flanking DNMTS sessions (**Fig. S4C**).

Are the ‘Sample’ and ‘Choice’ events of task context parsed by the patterned firing of the assembly member units? Rhythmic 4-5Hz co-modulation between dCA1-mPFC unit pairs was prominent only during the 4s preceding sample lever presses (**Fig. 4D**, **Fig. S4D**). This dynamic signature of dCA1-mPFC assembly activity therefore timestamped the DNMTS sample phase, and is consistent with evidence that optogenetic silencing of hippocampal-prefrontal interactions is particularly disruptive during the sample phase of delayed responding on a T-maze^64^.

In contrast, cross-correlations between dCA1-mPFC assembly member pairs preceding choice lever presses were not 4-5Hz modulated, but tended to reflect mPFC spiking leading dCA1 spiking on a longer, 400ms timescale (**Fig. 4D**, bottom right panel), consistent with previous studies of decision-making^60,70,71^. Contextdependent shifts in signal 4-5Hz modulation and flow direction between hippocampus and frontal cortex can therefore serve to disambiguate the transition from sample to choice phases of DNMTS.

We further explored this hypothesis by examining when individual assemblies were most active during sample or choice events. **Fig. 4E** summarizes the activity profiles of within-area (gold) and inter-area (purple) cell assemblies during the task, demonstrating comparable activation during sample presentation but diverging activation levels during delay and choice events. We restricted the assemblies to those whose factor scores showed significant activity (bootstrapped p<0.05 compared to assemblies composed from time-shuffled neural spike trains, **Fig. 4B**) in the 4s preceding lever presses, and split them into either significantly “sample-active” or “choice-active” based on the time of their strongest average activations (**Fig. 4F**, **Fig. S4D**).

A significant majority of within-dCA1 and dCA1-mPFC assemblies were sample-active (dCA1: 13 Sample-active vs. 5 choice-active; dCA1-mPFC: 20 Sample-active vs. 6 choice-active detected assemblies, one-sample t-test vs. an even Sample/Choice split for within-animal averages; mPFC: T(6)=-2.61, p=0.08, dCA1: T(6)=-2.53, p=0.13, dCA1-mPFC: T(6)=-6.72, p<0.01, Fig. 4F). In contrast, both classes of sample and choice activity were approximately equally represented in within-mPFC assemblies (9 Sample-active vs. 10 choice-active detected assemblies). This was explained neither by skewed unit counts across areas in our recordings nor over-representation of one area’s contribution to the FA assembly models (**Fig. S4E,F**). Finally, restricting analysis of rhythmic dCA1-mPFC unit pair interactions to sample-active assemblies amplified the 4-5Hz rhythmic coordination (**Fig. 4G**). The coherent modulation of spike trains by this fingerprint “Sample” rhythm was attenuated between parallel but independent dCA1-mPFC assemblies, and was absent in the activities of Sample assemblies during the choice-preparatory period.

These assembly analyses reveal sparse subsets of single units that cohere task-dependently into assemblies spanning dCA1 and mPFC, and are hallmarked by a physiological signature of 4-5Hz coordination. As such, the inter-area cell assemblies are uniquely positioned to orchestrate dCA1-mPFC interactions and information transfer.

Having uncovered dissociations between coding of dCA1 and mPFC populations, and task phase-dependent activation of 4-5Hz modulated dCA1-mPFC joint assemblies, we sought to map the activities of these subpopulations of neurons onto component processes of working memory underlying DNMTS task performance. Times at which the intra-CA1, intra-mPFC and dCA1-mPFC classes of assembly-participating units provided significant cue encoding during the task showed no clear segregation at the single cell level (**Fig. 5A**). However, multivariate population decoding from units belonging to these different assembly types demonstrated dynamic, fluctuating contributions of each neuronal class during the DNMTS task (**Fig. 5B**, **S5**). For example, mPFC inter-area assembly members (purple traces in **Fig. 5B**, **S5**) showed strong sample representation, while non-members (i.e. mPFC units less correlated with either dCA1 or other mPFC units) showed strong encoding during the delay.

**Figure 5:**
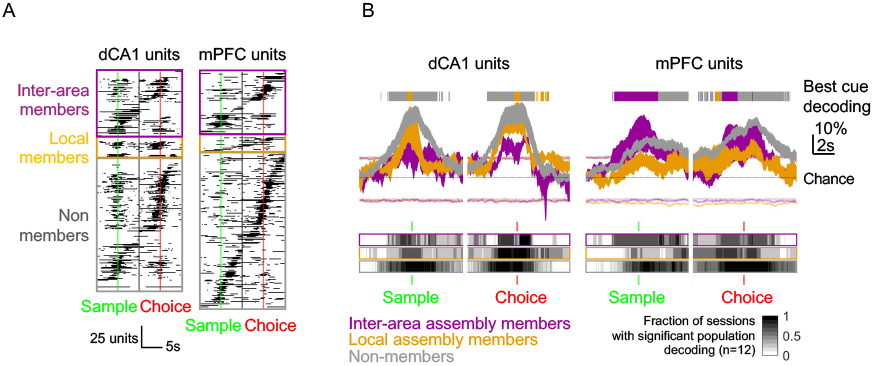
Assembly membership-dependent differences in the statistics of cue encoding during the DNMTS task are not visible at the single neuron level but orchestrate dynamic contributions by neural populations. **A** Discrimination of cue location by firing rates of individual dCA1 and mPFC single units, sorted by assembly membership type and time of peak discrimination. Black bars indicate time of significant decoding (t-test, Bonferroni-p<0.05, bootstrapped confidence limit). **B** Population-level discrimination of cue location by firing rates differs by class of assembly membership. Curves show cross-validated multivariate decoding for units participating in different assembly types (mean±SEM across recording sessions). Coloured lines indicate 5%, 95% bootstrap CIs of decoding from shuffled trial labels (mean across session shown). Below: Grey shading indicates fraction of recording sessions with decoding exceeding 95% CI at each time-point. Above: best performing group providing significant decoding from >60% sessions, winner-take-all.

These analyses demonstrate a dynamic modulation in the representation of cue information by cells forming assemblies that cannot be detected in the activities of individual member neurons. In particular, changes in the pattern of interaction across brain areas hallmarks parsing of the cognitive contexts of encoding, maintenance and recall lever presses during the task.

### Incorrect choices are associated with intact sample encoding in hippocampus but impaired transfer of cue information to mPFC and collapse of intra-mPFC dynamics

The distributed and dynamic mechanisms supporting successful completion of the DNMTS task offer multiple potential vulnerabilities to disruption, culminating in erroneous choices. We therefore quantified which features of coordinated population activity were altered on error trials.

The average activation profiles of within-dCA1 or sample-active within-mPFC assemblies were unaffected on error trials (**Fig. 6A**, top), whereas choice-active, within-mPFC assembly activation was significantly weakened in the period leading up to the choice lever press (gold vs. red traces in **Fig. 6A**, third row). No differences were observed in average activation strengths of inter-area dCA1-mPFC assemblies preceding the incorrect Choice lever presses, but activation was significantly stronger immediately afterwards (purple vs. red traces in **Fig. 6A**, bottom), suggesting coordinated inter-area firing in error feedback signalling, as has been reported for this pathway in primates^61^.

**Figure 6:**
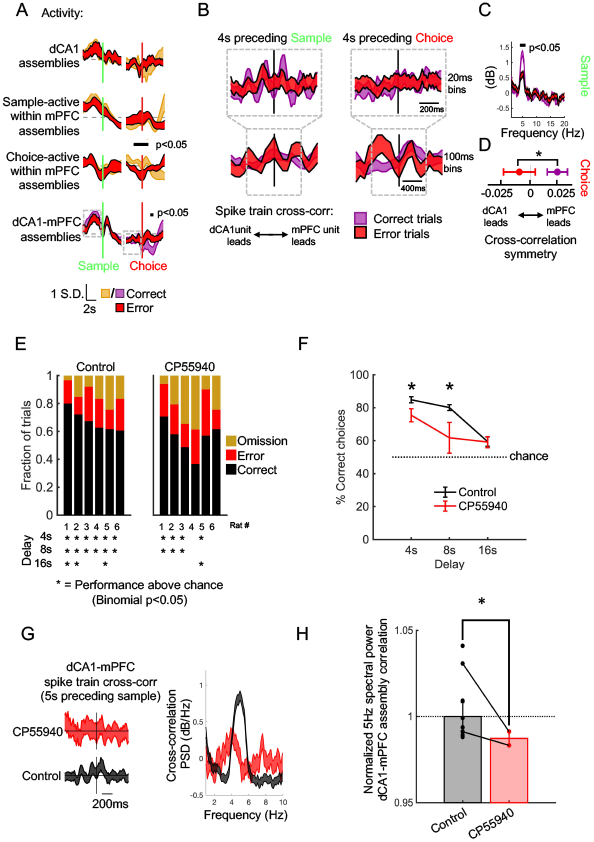
Incorrect choices are associated with intact sample encoding by dCA1 networks but reduced 4Hz dCA1-mPFC assembly synchrony, leading to a collapse of intra-mPFC dynamics and impaired dCA1-mPFC synchronization on recall. **A** Average Z-scored activity of different classes of within- and inter-area cell assemblies on correct (gold/purple) and error (red) trials. Mean±SEM across all pairs shown. Grey dashed lines indicate mean activity. Black bars: times of significant changes on error trials (Bonferroni-corrected bootstrap p<0.05, permutation test). Grey boxes indicate regions used for spike train cross-correlations in C,D. **B** Inter-area correlation between dCA1-mPFC cell pairs from the same assembly on correct (purple) and error (red) trials. Details as Fig. 4D, average of 251 pairs from 10 sessions shown. **C** Power spectrum of 20ms-binned cross-correlations in 4s preceding Sample in B, black bars indicate frequencies with significant change in power between correct and error trials (Bonferroni-corrected bootstrap p<0.05, permutation test). **D** Balance of dCA1-to mPFC driven correlation in 100ms-binned spike times from the 4s preceding Choice was significantly reversed on error trials. **E** Rats administered CP55940 performed worse in the DNMTS task. Bar indicates the fraction of trials of each outcome for each individual rat, sorted from left by best-performing animal on Control sessions. Asterisks below indicate whether the rats performed at- or above-chance for each delay length. **F** Group data from N=6 rats demonstrating significant impairment in task performance on 4s and 8s delay trials on CP55940 recording sessions. **G** Rhythmic 4-5Hz spike-train cross-correlations for within-assembly dCA1-mPFC unit pairs during cue sampling was strongly attenuated in rats administered CP55940. Left: rate-normalized cross-correlations (method as Fig. 4D). Right: power spectra of cross-correlograms. **H** Strength of rhythmic 4-5Hz dCA1-mPFC correlation between unit pairs (average power in spectrum between 3.5-5.5Hz), normalized by group mean on control days. Dots are individual animals, black lines link animals between control and CP55940 sessions.

Because, on correct trials, cue encoding by dCA1 -mPFC assemblies showed two distinct modes of correlated interaction during Sample and Choice periods (**Fig. 4E-G**), we wondered whether the rhythmic firing before Sample and mPFC-driven correlations before Choice times (Grey boxes in **Fig. 6B**) would be specifically affected on errors (**Fig. 6B**). Indeed, we observed significantly weaker 5Hz correlated firing between interarea pairs from the same assembly during the Sample-preparatory period (**Fig. 6C**), and a reverse in lag of peak correlation, such that dCA1 led mPFC firing in the Choice-preparatory period on error trials instead of mPFC leading dCA1 as in correct trials (**Fig. 6D**, t(250)=2.21, p=0.0274, paired t-test).

Given these disrupted rhythmic and directional interactions during unforced errors, we tested whether pharmacological impairment of working memory performance also associated with changes in CA1-mPFC 4-5Hz synchrony. On separate recording days, we systemically injected rats with the synthetic cannabinoid CP44940, previously reported to impair working memory performance^72^. We observed strong effects of the drug on animals’ performance in the DNMTS task (**Fig. 6E,F**). Compared to control sessions in which all six rats performed above chance on four- and eight-second delay trials (asterisks in **Fig. 6E**), CP55940 reduced performance to chance in half of the animals. We observed significant Drug- and Delay-dependent effects and a Drug x Delay interaction on performance (Fig. 6F, repeated-measures two-way ANOVA; Drug: F(1,5)=915, p<0.001, Delay: F(1,5)=1850, p<0.001, DrugXDelay: F(1,5)=899, p<0.001). Post-hoc tests (Tukey-Kramer p<0.05) indicated that CP55940 significantly reduced performance on the shortest, 4s and 8s delays trials. No changes in trial omission rates (Normalized to mean Control rates: Control: 100±20%, CP55940: 140±30% F(2,21)=0.74,p=0.49) or systematic effects on behavioural latencies (data not shown) were observed across different conditions, suggesting that the drug did not affect the animals’ engagement in the task.

We examined CP55940’s effects on rhythmic cross-correlations between dCA1-mPFC cell assembly member neurons during sample encoding (**Fig. 6G**). Compared to control sessions, the magnitude of 4-5Hz power in spike train cross-correlations when considering all within-assembly pairs was significantly reduced under CP55940 (**Fig. 6H**, Z=3.6, p=0.0032, n=252,78 dCA1-mPFC unit pairs from N=6,2 animals for Control, CP55940 sessions, respectively). Because dCA1-mPFC cell assemblies were detected in only two CP55940 sessions (although this did not represent a significant reduction in the number of dCA1-mPFC assemblies from control conditions: fraction of sessions with detected assemblies Control: 3/6, CP55940: 2/6, *χ*^2^(1) = 0.343,p=0.560, similar results for numbers of parallel detected assemblies: **Fig. S6A**), we also considered the distributions of peak 5Hz spike train correlations between all possible dCA1-mPFC unit pairs (**Fig. S6B**). The strongest modulated dCA1-mPFC pairs demonstrated a significant reduction under CP55940 (Komogorov-Smirnov test, D = 0.0047, p<0.001, n = 11686, 3212 pairs from N= 6,6 sessions, respectively). This is consistent with the observation that 4-5Hz correlated modulations are strongest between inter-area cell-assembly member pairs in control conditions during cue-sampling (**Fig. 4**) and that these are affected by cannabinoid administration.

Thus, as with naturally occurring unforced errors in DNMTS performance, biasing animals’ behaviour with psychoactive compounds known to affect working memory also implicates rhythmic hippocampal-prefrontal network population coordination during working memory loading in later incorrect choice-making.

## Discussion

The temporal structure of the operant DNMTS task disambiguates mnemonic and decision-making processes, which overlap in space and time during the maze-based tasks commonly employed in rodent studies of working memory. Simultaneous dCA1-mPFC recordings, allied with the temporal structure of the operant DNMTS task, unveiled several principal features of dCA1 and mPFC information processing. Encoding of trial-specific cue information during Sample is strongest in dCA1 and integrated into mPFC processing by virtue of joint dCA1-mPFC assemblies, bound by a common 4-5Hz rhythmic modulation. Separable mPFC populations then maintain sample information through sequential delay-spanning dynamics; on error trials this coding degrades, despite accurate encoding in dCA1. Finally, dCA1 and mPFC concurrently encode choice information, led by mPFC and no longer contingent upon the rhythmic assemblies recruited during Sample.

These differential dCA1 and mPFC coding patterns corroborate previous lesion studies in rodents^73–75^, population recordings in non-human primates^62^ and human imaging studies highlighting hippocampal activation during encoding of working memory^76–79^. However, we also resolved a physiologically distinct subset of dCA1 and mPFC neurons critical for trial-specific loading of cue identity into working memory, coactive as inter-regional assemblies with sub-50ms timescales.

Activation of dCA1-mPFC assemblies occurred during the DNMTS sample phase analogous to the “sample” run on a T-maze, when optogenetic silencing of CA1 projections to mPFC preferentially impairs task performance^64^. These coordinated activities reflect hippocampal-prefrontal interactions during loading of working memory, with the tendency of dCA1 encoding to precede mPFC encoding consistent with hippocampal-to-prefrontal anatomy^80–82^ and functional connectivity^60^. Conversely, delay-coding mPFC subpopulations may partner with mediodorsal thalamus, since optogenetic disruption of mPFC activity^83^ or silencing mPFC input from mediodorsal thalamus during the delay phase of short-term memory tasks impairs maintenance of information^24^. Based on recent evidence that individual mPFC pyramidal neurons receive convergent input from both ventral CA1 and mediodorsal thalamus^84^, which neurons are recruited during sample vs. delay may therefore rather reflect dynamic configuration of assemblies via interneuron-mediated tuning of excitability and synaptic plasticity^85^.

Rhythmic coordination of dCA1-mPFC assembly members is reminiscent of a 4Hz rhythm reported during working memory processing in rats^86^, coordinating activity across hippocampus, ventral tegmental area and mPFC during delayed responding on a T-maze. In the DNMTS task, we pinpoint its emergence during the sample phase and specificity to the units forming inter-regional dCA1-mPFC assemblies, evident both in spike cross-correlograms and in their intrinsic auto-correlated firing. In contrast, 4-5Hz dCA1-mPFC assembly modulation was absent around memory-guided DNMTS choice lever presses, showing that hippocampal-prefrontal dynamics are reconfigured from sample to choice, reflecting mPFC-led control of memory retrieval^55^.

The cross-temporal coding analyses in Figure 2 evidence sustained coding despite transient activities of individual mPFC neurons. This delay coding was neither evident in dCA1, nor associated with sustained activation of dCA1 or mPFC synchronous assemblies; it is most likely, therefore, to derive from sequential activation of mPFC units and/or assemblies^47^, or potentially from assemblies more widely distributed across PFC than our tetrode recordings could sample. Whatever its basis, sustained mPFC coding during the DNMTS delay phase collapsed during errors, and on 16s delay trials during sessions in which rats performed at these trials at chance levels. In fact, rats may resort to guessing on both correct and error trials during chance performance sessions, though consistent nose-poke and lever-press latencies across trials and sessions indicate that motivation and certainty/expectancy remained stable.

On error trials, the dCA1 population code maintains a faithful representation of cue identity, and activation strengths of cell assemblies linking dCA1 and mPFC are comparable to correct trials during both Sample and Choice epochs. However, transient Sample-specific rhythmic interactions are weaker, as are the network dynamics in mPFC that maintain cue identity encoding during the delay, preventing the formation of a stable population code for working memory. Although with the current dataset we cannot dissect causal contributions of individual signatures of errors, such sequential events could lead to the disorganised activation of choice dCA1-mPFC cell assemblies we observe preceding incorrect choices.

Disrupted hippocampal-prefrontal connectivity impact working memory performance^58,64–66,72,73,87^ and is implicated in the pathophysiology of schizophrenia^88,89^. Here, we systematically disrupted rhythmic dCA1-mPFC sample assembly activation through systemic activation of CB1 receptors by CP55940. Correspondingly, delay-dependent task error-making was exacerbated, potentially via combined effects on cortical glutamatergic transmission^90^ and hippocampal ensembles^91^.

Thus, whether arising spontaneously or induced by CP55940, interruptions to population coordination either during initiation of dCA1-mPFC encoding during the sample phase or intra-mPFC processing during the maintenance phase associate with impaired delay-dependant performance on error trials, leading to incorrect choices. In conclusion (schematized in **Fig. 7**), our data confirm why both mPFC and dCA1 – as well as intact connectivity between them – have been ascribed crucial roles in spatial working memory. The temporally defined set of cognitive steps timestamped by rhythmic assembly dynamics establishes a framework that can now be tested in combination with circuit tracing and/or imaging strategies relating assembly configurations to the connectivity of participating neurons and their neuromodulation.

**Figure 7:**
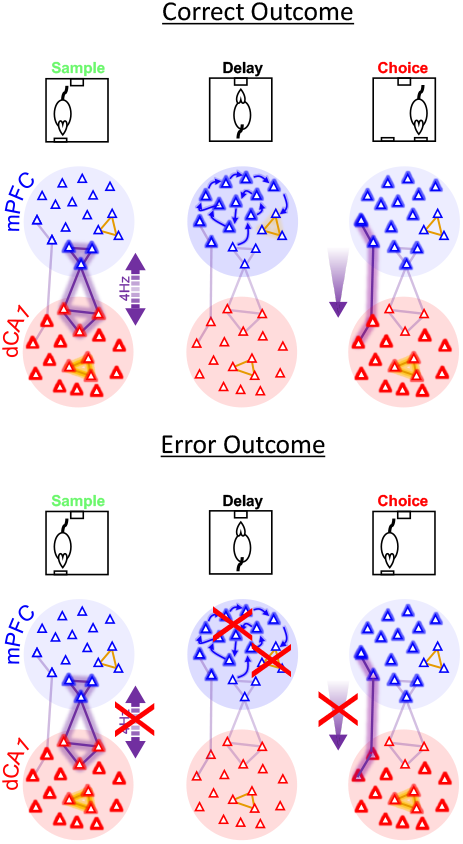
Schematic of main findings Lines indicate cell assembly motifs within and between brain regions, triangle outline width represents strength of cue encoding by individual neurons and cell assemblies of each type, for different phases of the task. Arrows indicate task information flow; either rhythmic (dotted) or directional (shaded). Red crosses highlight physiological mechanisms of information encoding observed to collapse on error trials.

## Methods

### Electrode implantation

All procedures were conducted in accordance with the UK Animals Scientific Procedures Act (1986) and with the approval of the University of Bristol Ethics Committee. Eight adult (300–400g) male Long–Evans rats (Harlan UK) were implanted with 16 extracellular tetrode recording electrodes: 8 over right medial prefrontal cortex (+3.2 mm, +0.6 mm from bregma) and 8 over the right dorsal hippocampus (−4.0 mm, +2.5 mm from bregma) under sodium pentobarbital recovery anaesthesia. During 7–12 days following surgery the independently moveable tetrodes were lowered into prelimbic cortex (~2–3 mm ventral) and the principal cell layer of the dCA1 ^92^, guided by the characteristic burst mode of single-unit firing and the presence of large-amplitude sharp-wave ripple events in the local field potential. Extracellular action potentials (sampled at 32 kHz and filtered between 0.6–6 kHz) together with local field potentials (sampled at 2 kHz and filtered between 0.1–475 Hz) were recorded differentially (Digital Lynx, Neuralynx) using local references, which were targeted to superficial prefrontal cortex and the white matter overlying the hippocampus. Two screws placed in the skull overlying the cerebellum were used as ground connections. Final tetrode tip positions were verified histologically by identifying sites of electrolytic lesions in 50um stained sections of formaldehyde-perfused brain.

### Behavioral training

Subjects were food-restricted to no less than 85% of their free-feeding weight and trained in a DNMTS operant task (**Fig. 1**). We used an operant chamber (Med-Associates, Vermont, USA), which consisted of two retractable levers facing a food pellet dispenser on the opposite wall, with a cue light above each component and a tone generator placed above the pellet dispenser. Side and top walls of the chamber were transparent to enable view of distal spatial cues in the recording room. Metal components of the chamber were grounded to the amplifier to electrically shield the recordings, which were carried to the data acquisition system via tethers suspended through a hole in the centre of the box ceiling. The task was programmed and operated in K-Limbic software (D. Fuller, Conclusive Marketing Ltd.) on a separate computer. Subjects were initially conditioned to press a lever to obtain pellet reward before being trained in DNMTS task with pseudo-random delays (random combination of equal number of target left and right lever trails at each delay arranged into blocks of 10 trials) of 4,8, 16s. Error and missed trials were followed by all cue lights off for an extra 10s of inter-trial interval. There were 150 trials in each session (50 x 3 delays). Sessions with less than 67% of trials completed were excluded from further analysis.

### Single unit clustering

Single units were isolated off-line using automated clustering software (KlustaKwik 1.7; K. Harris), followed by verification and manual refinement in MClust 3.5 (A.D. Redish); unit inclusion criteria were set to isolation distance >10.0 and L-ratio <0.35, with <2% of spikes within 2ms inter-spike interval (Suppl. Fig. 1). Putative pyramidal cells were classified based on the spike width, waveform and mean firing rate. A total of 156 (min. 115, max. 194) putative principal cells in dCA1 (mean of 34 units per subject) and 168 (min. 152, max. 201) putative pyramidal cells in mPFC (mean of 33 units per subject) were isolated in each recording session.

### Statistical analysis methods

Where appropriate following normality testing (Kolmogorov-Smirnov test, p>0.05), parametric statistical comparisons were performed. Unless otherwise specified, results are quoted as Mean ± Standard Error of the Mean (SEM). To equalize statistical power on multivariate statistical analyses, all comparisons across delay lengths and different recording sessions were calculated on repeated jack-knife draws of random subsets of matched numbers of trials. Similarly, non-parametric bootstrapping was performed through calculating statistics on distributions of shuffled data (e.g. for decoding analyses described below, by randomly permuting trial labels 1000 times). Results were considered significant if the observed value exceeded the 95^th^ percentile of the bootstrap distribution. Two bootstrap resampled distributions were considered significantly different if their <5% and >95% tails did not overlap. Where two time series were compared (e.g. **Fig. 1F,G**), bootstrapped p–values were adjusted using Bonferroni correction for number of time bins.

### Spike train analysis

Only units with an average firing rate of at least 0.1 Hz were included in all subsequent analyses, but no further selection of units was performed. All decoding analyses, to be described further below, were performed on kernel density estimates of the instantaneous spiking rate. Separate kernel density estimates (KDE) for each unit ***i*** were obtained by convolving spike trains with Gaussian functions (‘kernels’), where the optimal kernel width σ^2^ was determined through unbiased cross-validation ^93^. For Gaussian kernels, closed-form expressions for the unbiased cross-validation error (CVE) can be obtained, and numerical iteration of the CVE procedure is not necessary ^93^. Loosely, one may think of the unbiased cross-validation procedure as leaving out each spike in turn, and evaluating the likelihood of its actual position from the spike density estimate obtained based on all other spikes in the series. Thus, the optimal bandwidth estimated will depend on predictable temporal structure in the spike trains, not just their rate (see also Shimazaki and Shinomoto, 2010). KDEs provide a statistically more robust (less variable) estimate of the true underlying spike density, compared to e.g. histograms or binarized spike series, but decoding results did not crucially depend on this pre-processing step.

Single units were considered significantly sensitive to behavioural events if their normalized (z-scored) firing rate deflection in a 2 second window after the event exceeded ±3x the standard deviation of the baseline firing rate (500ms window preceding the event).

### Neural Decoding

For single unit decoding (e.g. **Fig. 2A**), for each time bin *m* and unit *i* single unit rates *v_im_* were collected into two sets according to whether *m*∈C1 or *m*∈C2, the two sets of time bins associated with one (C1) or the other (C2) cue stimulus. The common t-statistic (as also employed in Student’s two-sample t-test) is a measure of discrimination among these two sets, as it divides the difference in means by the pooled standard deviation (c.f. Durstewitz et al., 2010). For the average number of trials collected here (~54), values of approximately t > 1.67 would indicate significant discrimination at the p<0.05 level.

Leave-one-out cross-validation analysis (e.g. **Fig. 1F**) was performed for the multivariate linear discriminant classifiers used for decoding (e.g. Hastie et al., 2011), with regularized covariance matrix as specified below. This used, for each time bin *t* the two sets of population vectors associated with the two stimulus classes (see above), with one population vector (and thus trial) left out from the fitting. Prediction performance was evaluated on the left-out trial, and this was repeated for each trial in turn, yielding the cross-validation error (CVE) as the relative number of incorrectly classified (out-of-sample) prediction trials. For testing differences in CVE between mPFC and dCA1 populations, for each data set the number of mPFC and dCA1 units (variables) used for decoding was exactly equalized to rule out any potential confounds due to population size. This was done by fixing the number K of units used to the smaller of the two populations, mPFC or dCA1, and then randomly drawing K units with replacement from the larger of the two populations 10 times and averaging the obtained CVE values. Differences in relative proportions of correct cue predictions, CP = 1-CVE ∈ [0, 1], between mPFC and dCA1 were statistically tested by averaging CP across all 12 data sets and using the beta distribution. Specifically, at each time point *t* the smaller of the two values CP_mPFC_ and CP_dCA1_ was used for the reference distribution, and it was checked whether the larger of the two significantly (p<0.05) escaped this reference distribution given the average number of trials recorded.

To compare decoding performance on correct and error trials (e.g. **Fig. 1G**), classifiers trained on correct trials were additionally challenged to predict the cue identity of error trials in a similar manner as above.

To evaluate the stability of the population code for cue location during the delay period we used cross-temporal decoding methods inspired by Stokes et al. (2013), and Parthasarathy et al. (2017). Briefly, we performed leave-one-out cross-validated decoding of cue location from multi-single unit firing rates as described above using separate training and testing sets offset by sequential 50ms increments. The performance of the decoder at each combination of [train,test] time points is thus the percentage of test trials in which the decoder could correctly identify the cue location when trained using trials at a given time point.

To equalize size of training sets across combinations of delay lengths and recording sessions, cross-temporal decoding analysis was performed on random draws of eight trials from each of left and right cue conditions, repeated 500 times. Thus the total cross-validation size for each decoder was 5000 random resamples per [train,test] time combination. Mean performance across runs is reported in the colormaps shown in (e.g.) **Fig. 2B**. Significant decoding at each [train,test] point was calculated against distributions (p<0.05) from 1000 bootstrap draws created by shuffling labels. For visualization, a 250ms Gaussian smoothing kernel was applied after significance testing across both training and testing dimensions.

To further corroborate the classification results, we also used a parametric test statistic (**Fig. S5**): Vectors **v**_m_=(**v**_1m_…**V**_pm_)^⊤^ of all unit activities were collected into two sets corresponding to stimulus conditions as above, and contrasted by Hotelling’s T^2^ statistic, a multivariate generalization of the univariate two-sample t-statistic which relates differences in cue specific mean vectors to the pooled covariance matrix of the data ^67^. Hotelling’s T^2^, scaled by the appropriate degrees of freedom, is approximately F-distributed which can be used to construct parametric confidence bands.

In the present case, the number of recorded units often reaches or even exceeds the number of trials, causing singularity and over-fitting issues with the covariance matrix. One standard statistical remedy is regularization, where the covariance matrix Σ is moved toward the identity, Σ_*reg*_ = Σ + *λ**I***, with regularization parameter *λ* (set to 0.05 here, without any attempt to optimize this parameter; Hastie et al., 2011).

### FA model for cell assembly detection

Extraction of cell assemblies was based on Factor Analysis^68^. This model-based statistical tool is designed to extract *correlations* between variables. It assumes that observation vectors **v_i,t_** are given by a (linear) mixing of uncorrelated latent variables (factors) **z_i,t_**, plus common mean **μ_i_** and measurement noise **ε_i,t_**,

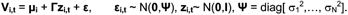

Parameters are commonly estimated through maximum-likelihood. Unlike principal component analysis (PCA) which detects variance-maximizing directions, FA attempts to capture all the *correlations* among the observed variables through the mixing of uncorrelated factors (e.g. Krzanowski, 2000). It is thus more appropriate for assembly detection than PCA, as has been demonstrated before^69,97^.

Inputs to FA were the kernel density estimated instantaneous firing rate vectors **C**_m_=(*c*i_m_) which collected spike rates cm for each unit *i* at time *m* binned at 50ms in columns, excised from time periods from cue presentation −5s to choice lever press +5s, combined from all trials.

Each of the 12 recorded data sets was treated separately, with simultaneously recorded mPFC only, dCA1 only, or concatenated mPFC and dCA1 units submitted for assembly analysis.

The likelihood-ratio statistic for FA models of increasing complexity (i.e. increasing number of factors) in conjunction with confidence bands obtained from trial-shuffled data can be used to determine the number of putative assemblies (i.e. significant factors) present (**Fig. 3B**). While, in principle, likelihood-ratio based parametric F-scores could be used to determine whether adding another factor to the model still significantly improves the fit, here we relied on Ho distributions generated from trial permutation bootstraps to account for the time series (and thus potentially dependent) nature of the data. Specifically, if **c**_i_,^(k)^ = (c_i1_^(k)^…c_iM_^(k)^) denotes the set of firing rates for unit *i* on trial *k,* for each unit separately the assignments of these sets to trials *k* were randomly shuffled. Thus, all autocorrelations and the firing rate structure across a trial were preserved for each unit *i* in the bootstrap data, while cross-dependencies between units were destroyed. These bootstrapped data sets (total of 500) were used both to determine the number of significant factors, i.e. those for which the LLR ranged within the 1% upper confidence limit of the bootstrap data, as well as significant factor loadings (the correlations of the units with the factors): Only units for which a *factor loading* exceeded 1% of the bootstrap range were assigned to the respective assembly. For each factor, the *factor score* (the value *z_lm_* on factor *l* in time bin *m*) quantifies the degree to which the respective assembly is activated. Local cell assemblies forming subsets of joint area assemblies were assumed as part of the larger assembly. Assembly detection by FA was confirmed using another method based on Independent Component Analysis ^98^. Assembly units were determined using IC weight threshold of 2.5 S.D. for every spike train in the analysis. As done for the FA based analysis, neuronal assemblies, defined as groups of three or more single units that consistently co-activated within a 50ms time window, were thus detected in dCA1, mPFC, and across dCA1 and mPFC. Despite this quite different methodological approach, sets of assemblies detected by ICA were highly similar to those detected by FA as quantified through the measure of overlap O=|A∩B|/|A∪B|∈ [0,1] between pairs of sets as defined further above. For each assembly set detected by ICA, the corresponding assembly set with highest similarity to it as detected by FA was first determined, and the average across O from all these ICA x FA pairs then calculated for each data set. Overall, across all data sets, there was an 84% average agreement between FA and ICA assemblies (**Fig. S3**).

### Pharmacology

CP55940 (Tocris) was diluted in a vehicle solution (10% ethanol, 10% cremophor, 80% saline) before the experimental session. Drug solutions were administered I.P. at the injection volume of 0.1ml/100g of animal body weight to a final concentration of 0.30 mgkg^-1^. Subjects were tested at 30min post-injection.

## Supplementary figures

**Figure S1:**
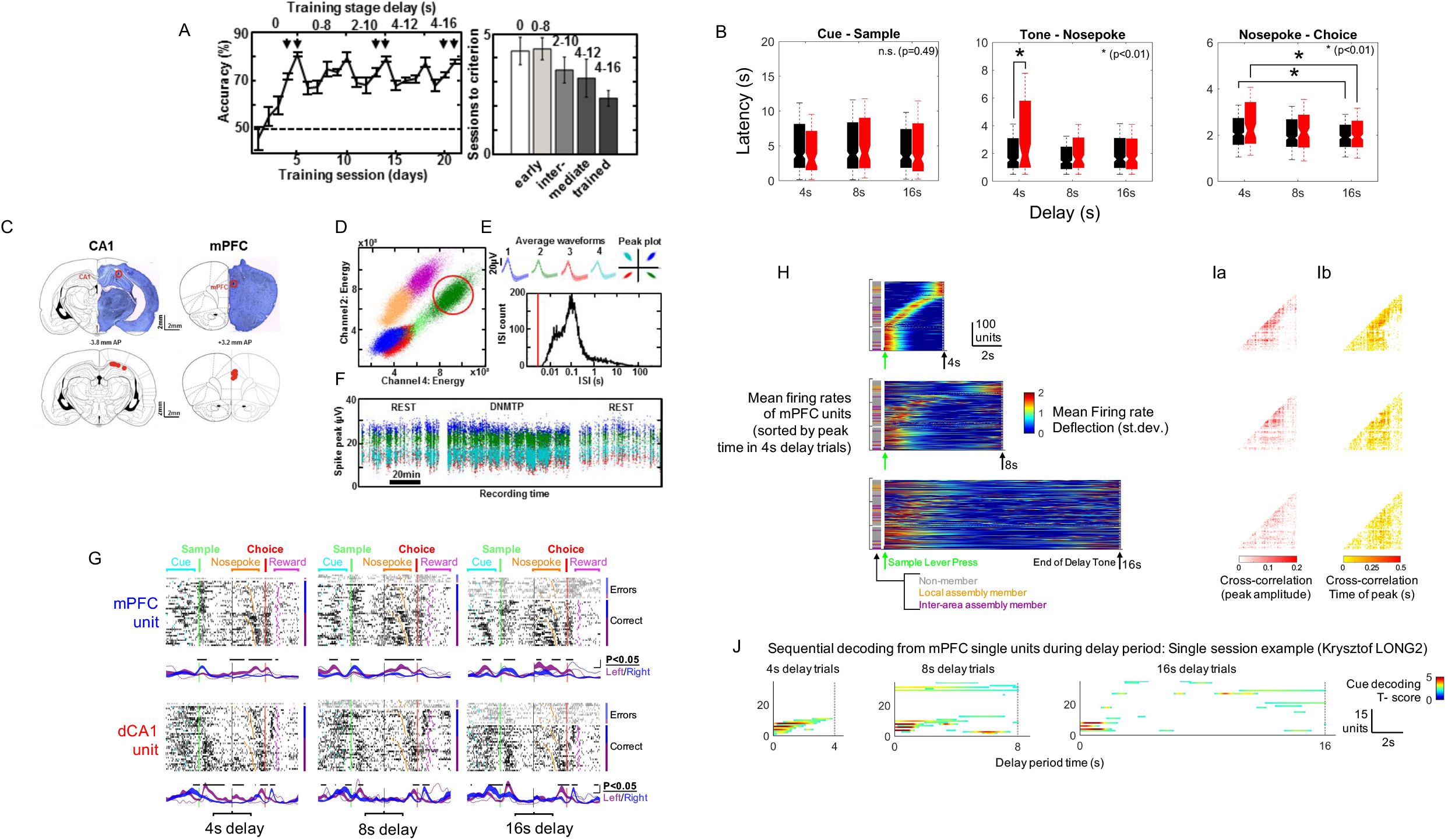
Details of training, recordings and single unit physiology during DNMTS task. **A** Behavioral performance of rats in DNMTS training is expressed as choice accuracy across subsequent sessions of the training (left) and as average number of sessions required to reach criterion performance at different stages of training: early (right). Arrows mark the two sessions used in the analysis of each stage (early, intermediate, trained). Data from the ”trained” days is analysed in this study. **B** Cue-Sample, Delay-Nosepoke, Nosepoke-Choice latencies averaged across animals and recording sessions. Asterisks at upper right corner indicate significant difference between conditions (Kruskal-Wallis ANOVA: *χ*^2^(5,1751)=3.69/16.29/16.53, p>0.05/p<0.01/p<0.01, respectively). Bars/asterisks indicate significant differences between specific combinations of delay and outcome conditions (Tukey-Kramer post-hoc test, p<0.05). **C** Coronal sections show example locations of tetrode recording sites (red circles mark the site of electrolytic lesions) in the prelimbic cortex (mPFC) and in the pyramidal cell layer of the CA1 subfield in dorsal hippocampus (CA1), matched to a corresponding rat brain atlas schematic (from Paxinos 2008). The lower panels summarize lesion sites across all six rats. **D** Extracellular action potential spikes recorded across an entire session were clustered into separate single units (colored dots), plotted here as waveform energy recorded on two channels of one mPFC tetrode. The properties of the cluster in red circle are presented in E and F. **E** Mean waveforms recorded on color-coded channels of the tetrode (top left) show consistent relative peak amplitudes (top right). Distribution of inter-spike intervals (ISI) below show no spikes detected in the <2ms refractory period. **F** Spike peak amplitudes of the same unit recorded on the color-coded four tetrode channels remain stable across the recording session. **G** Multi-trial firing raster from one example mPFC (top) and dCA1 (bottom) single units. Spike rasters with continuous firing rates aligned to the Sample and Choice lever presses +/-5s, with a variable portion of the delay period excised depending on delay length. Trials are sorted by correct and error outcomes (black and grey ticks) as well as left and right trial type, and finally by nose-poke latency. Solid areas indicate mean +/− SEM firing rates on correct trials, dotted lines show mean firing rate on error trials. Black bars above epochs show significant separation of Left/Right trial responses, from the trial-averaged firing rates (t-score, in 50ms non-overlapping increments, Bonferroni-corrected). **H** Mean firing rates of mPFC units (preferred cue direction, correct trials, all units across sessions combined) during the delay period sorted by time of peak firing on 4s delay trials, sort order maintained for 8,16s delay trials. Colored stripes on Left indicate assembly membership class (See Figures 3,4). **I** Firing rate correlation matrices (Ia: peak correlation and Ib: time-lag at peak) for data shown in H. Heat-map shows mean correlation across trials, note strong non-zero lagged correlations. **J** Sequential contributions of individual mPFC single units to maintaining population-level encoding of cue location during working memory delay. Single recording session shown. Units are shown sorted by center-of-mass of significant decoding (bootstrapped Bonferroni p<0.05) on 4s delay trials. Sort times maintained across longer delay lengths. Colourmap as for figure 1D.

**Figure S2:**
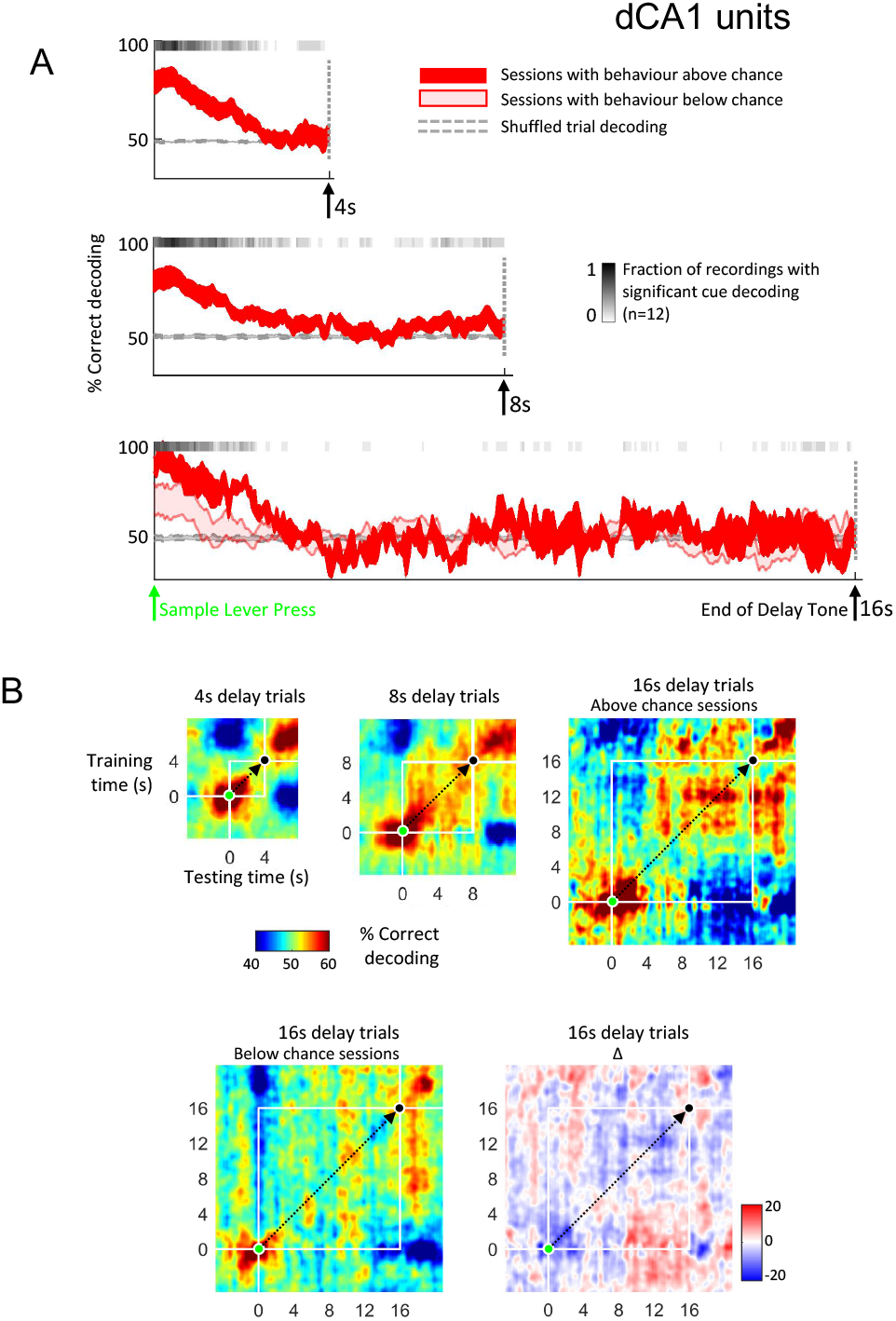
Transient cue encoding in dCA1 population lacks a stable code. **A-B**: Encoding of cue information during the delay. Legends as for figure 2 but decoding from populations of dCA1 single units.

**Figure S3:**
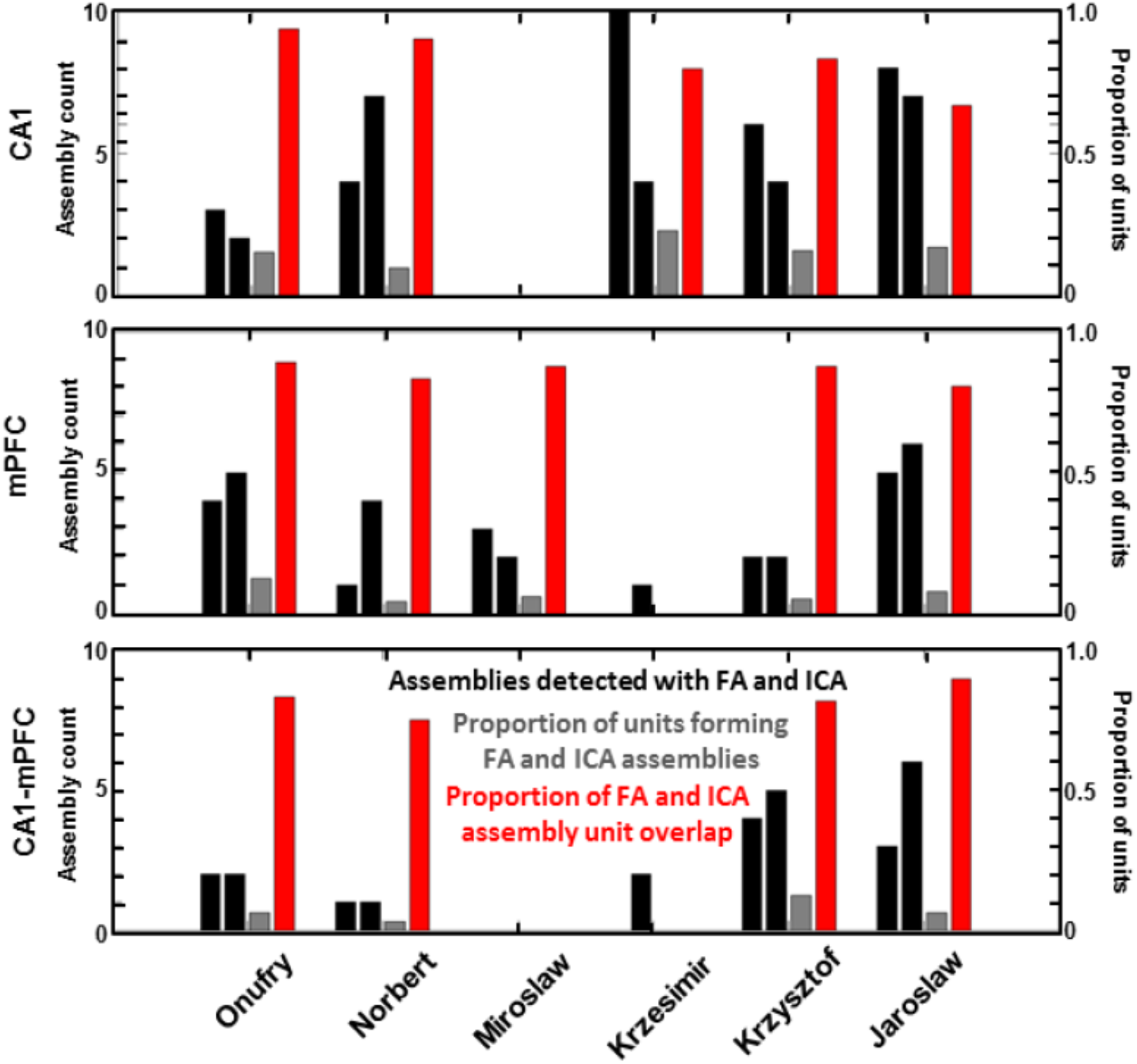
Validation of assemblies detected with the FA against PCA-ICA based methods. Total number of assemblies detected (black bars – the first for FA, second for ICA), proportion of units that participated in assemblies detected by both FA and ICA relative to all units recorded (grey), and proportion of unit overlap between matched FA and ICA assembly pairs (red) are summarized for the two sessions of each rat. A measure of overlap between a pair of assembly sets A and B was formally defined as O=|A⊣B|/|A∪B|∈ [0,1], i.e. the cardinality of the intersection divided by the cardinality of the union. On average there was 84% overlap between units detected by the FA and ICA methods, with similar numbers for total counts and unit proportions involved.

**Figure S4:**
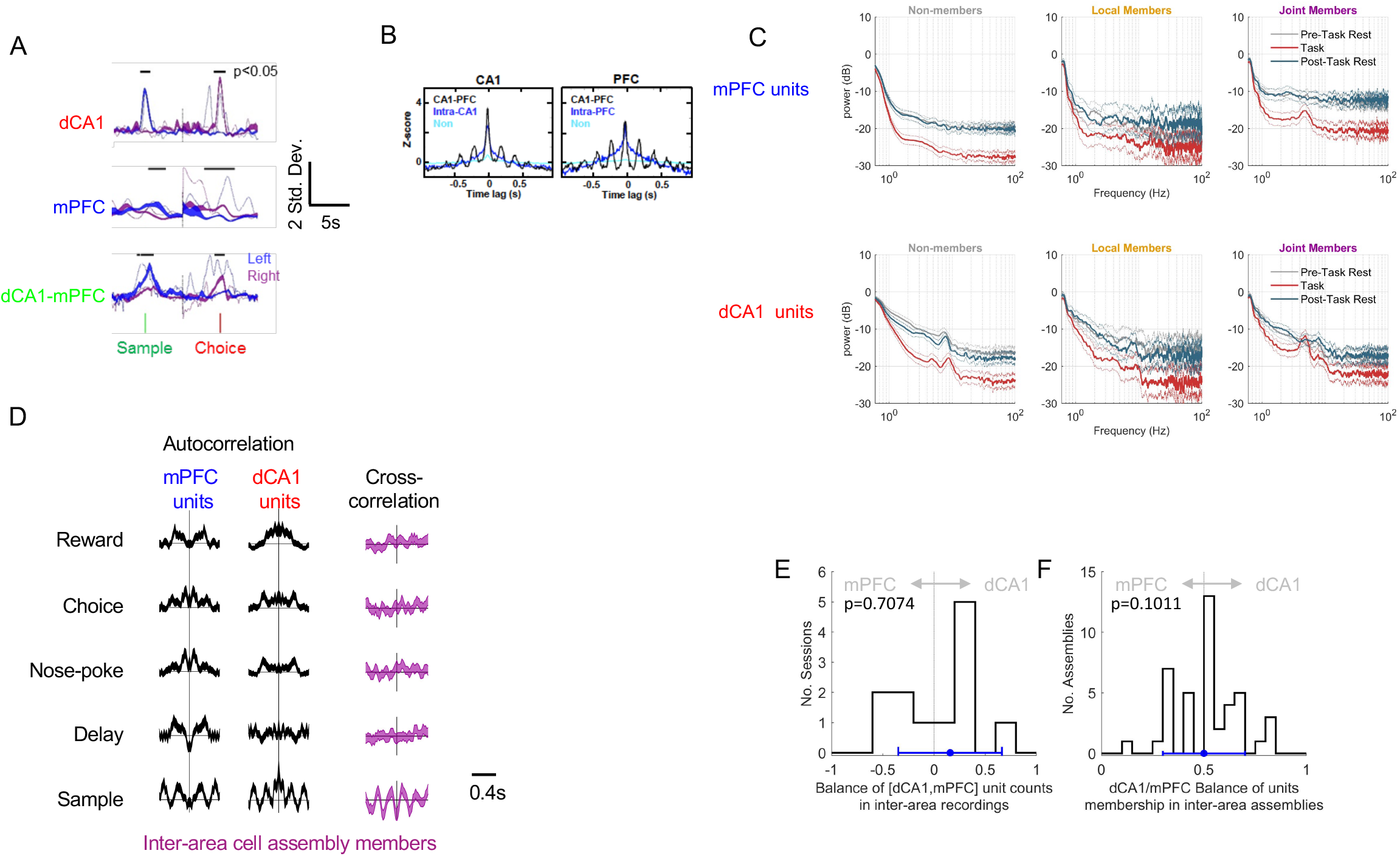
Further details regarding physiology of dCA1-mPFC cell assemblies. **A** Cell assemblies link single units conveying similar information. Example activities of one simultaneously recorded cell assembly from each of dCA1 and mPFC, and one inter-regional cell assembly, along with the firing activities of their member units. Shaded Blue/Purple lines indicate mean±SEM Factor Scores (cell assembly activity) on Left/Right trials, aligned to the lever press events. Thin lines represent mean firing rates of the member units of each assembly on Left/Right trials. Note the similar cue-selective activity and firing profiles of assemblies and member cells. Black bars indicate times when average Left and Right cue-related cell assembly activity differed between cues (p<0.05, bootstrap permutation test). **B** Averaged cross-correlograms of dCA1 (left) and mPFC (right) cell pairs drawn from either assemblies spanning the two regions (CA1-PFC, black), assemblies contained within one region (‘intra’ blue) or nonassembly neurons (cyan) from all recording sessions. Assembly pairs are more correlated than non-assembly pairs (black/blue vs. cyan), but 4-5Hz rhythmicity is specific to the CA1-mPFC co-assemblies (black vs. blue/cyan). **C** Power spectral densities of spike time autocorrelations for units of each area sorted by assembly membership category. Curves show mean±SEM spectra, Spikes are drawn from either the task period (red), or 1h pre- and post-task rest periods (grey, teal). For both dCA1 and mPFC units, the 4-5Hz oscillation was specific to the task period and, in mPFC, to cross-regional assembly members. **D** Units from both regions are equally represented in our recordings: Histogram across sessions of ratios between single unit counts in multi-area recordings [(#*units_mPFC_* – #*units*_*dCA*1_)/(#*units_mPFC_* + #*units*_*dCA*1_)]. Ratio distribution was not significantly different from a normal distribution (Kolmogorov-Smirnoff test). Blue symbols indicate median ± inter-quartile range. **E** Inter-area cell assemblies are equally contributed by dCA1 and mPFC single units. **F** Ratio distribution was not significantly different from a normal distribution with mean=0.5. (Kolmogorov-Smirnoff test). Blue symbols indicate median ± inter-quartile range.

**Figure S5:**
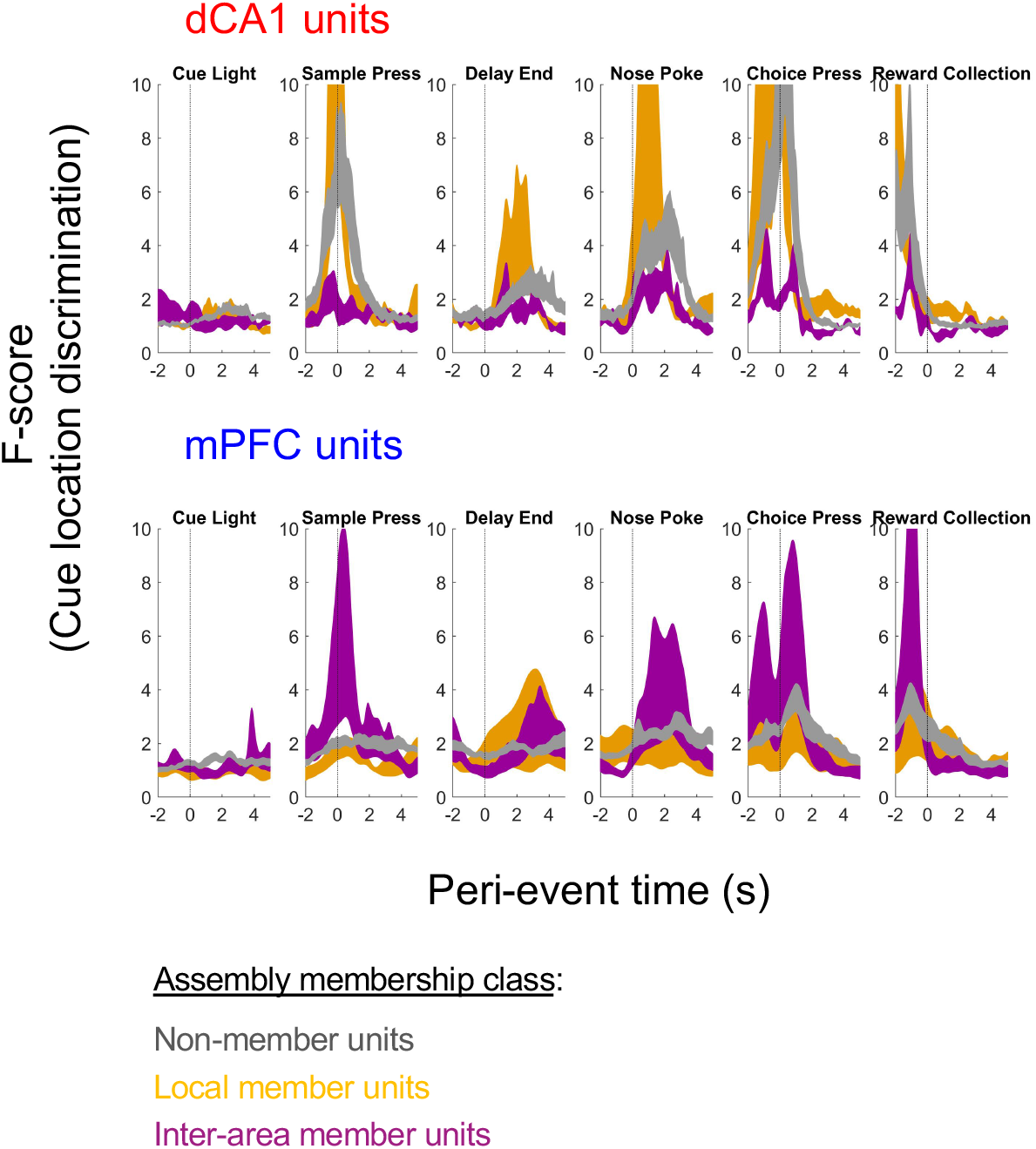
Evolution of cue discrimination during the DNMTS task by populations of dCA1 and mPFC units is determined by cell assembly membership participation. Time-aligned multivariate cue discrimination (regularized F-scores) of populations of each type of units for each event in the DNMTS task. Shaded regions indicate mean±SEM F-scores for units of each membership classification, from the 12 recording sessions (see methods for details).

**Figure S6:**
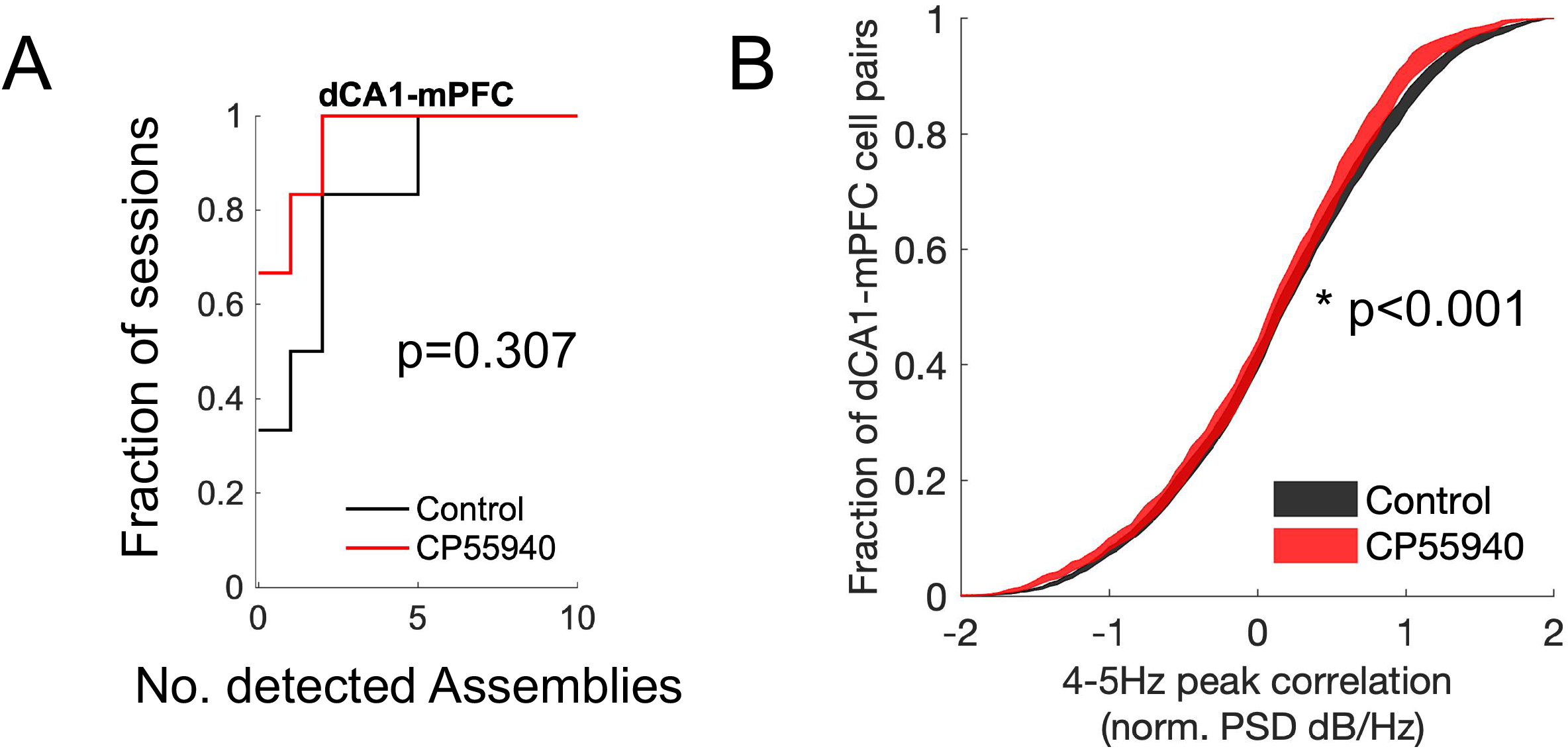
Further details of physiology of dCA1-mPFC cell assemblies in rats under influence of CP55940. **A** Numbers of detected parallel dCA1-mPFC cell assemblies in recordings of each condition (cumulative distributions). **B** Strength of 4-5Hz dCA1-mPFC spike train modulation during Sample phase (average power between 3.5–5.5Hz in spectrum of cross-correlation) for all inter-regional cell pairs, irrespective of cell assembly membership class. Shaded regions indicate mean±SEM of distributions across sessions (See main text for statistics).

## References

1. Baddeley, A. Working memory: theories, models, and controversies. Annu. Rev. Psychol. 63, 1–29 (2012).

2. Yu, J. Y. & Frank, L. M. Hippocampal-cortical interaction in decision making. Neurobiol. Learn. Mem. 117, 34–41 (2015).

3. Euston, D. R., Gruber, A. J. & McNaughton, B. L. The Role of Medial Prefrontal Cortex in Memory and Decision Making. Neuron 76, 1057–1070 (2012).

4. Fuster, J. M. Unit activity in prefrontal cortex during delayed-response performance: neuronal correlates of transient memory. J. Neurophysiol. 36, 61–78 (1973).

5. Funahashi, S., Bruce, C. J. & Goldman-Rakic, P. S. Mnemonic coding of visual space in the monkey’s dorsolateral prefrontal cortex. J. Neurophysiol. 61, 331–349 (1989).

6. Chafee, M. V. & Goldman-Rakic, P. S. Matching patterns of activity in primate prefrontal area 8a and parietal area 7ip neurons during a spatial working memory task. J. Neurophysiol. 79, 2919–2940 (1998).

7. Rainer, G., Asaad, W. F. & Miller, E. K. Selective representation of relevant information by neurons in the primate prefrontal cortex. Nature 393, 577–579 (1998).

8. Romo, R., Brody, C. D., Hernández, A. & Lemus, L. Neuronal correlates of parametric working memory in the prefrontal cortex. Nature 399, 470–473 (1999).

9. Hampson, R. E. & Deadwyler, S. A. Ensemble codes involving hippocampal neurons are at risk during delayed performance tests. Proc. Natl. Acad. Sci. U. S. A. 93, 13487–13493 (1996).

10. Hampson, R. E. & Deadwyler, S. A. Temporal firing characteristics and the strategic role of subicular neurons in short-term memory. Hippocampus 13, 529–541 (2003).

11. Hampson, R. E., Heyser, C. J. & Deadwyler, S. A. Hippocampal cell firing correlates of delayed-match-to-sample performance in the rat. Behav. Neurosci. 107, 715–739 (1993).

12. Deadwyler, S. A. & Hampson, R. E. Ensemble activity and behavior: What’s the code? Science (80-.). 270, 1316–1318 (1995).

13. Durstewitz, D., Vittoz, N. M., Floresco, S. B. & Seamans, J. K. Abrupt transitions between prefrontal neural ensemble states accompany behavioral transitions during rule learning. Neuron 66, 438–448 (2010).

14. Durstewitz, D. & Seamans, J. K. Beyond bistability: biophysics and temporal dynamics of working memory. Neuroscience 139, 119–33 (2006).

15. Falco, E. De et al. The Rat Medial Prefrontal Cortex Exhibits Flexible Neural Activity States during the Performance of an Odor Span Task. eNeuro 6, (2019).

16. Funahashi, S. Functions of delay-period activity in the prefrontal cortex and mnemonic scotomas revisited. Front. Syst. Neurosci. 9, 2 (2015).

17. Constantinidis, C. et al. Persistent Spiking Activity Underlies Working Memory. J. Neurosci. 38, 7020–7028 (2018).

18. Wang, X. J. Synaptic reverberation underlying mnemonic persistent activity. Trends Neurosci. 24, 455–63 (2001).

19. Zylberberg, J. & Strowbridge, B. W. Mechanisms of Persistent Activity in Cortical Circuits: Possible Neural Substrates for Working Memory. Annu. Rev. Neurosci. 40, 603–627 (2017).

20. Bouchacourt, F. & Buschman, T. J. A Flexible Model of Working Memory. Neuron 103, 147–160.e8 (2019).

21. Hart, E. & Huk, A. C. Recurrent circuit dynamics underlie persistent activity in the macaque frontoparietal network. Elife 9, 1–22 (2020).

22. Parnaudeau, S. et al. Inhibition of Mediodorsal Thalamus Disrupts Thalamofrontal Connectivity and Cognition. Neuron 77, 1151–1162 (2013).

23. Wimmer, R. D. et al. Thalamic control of sensory selection in divided attention. Nature 526, 705–709 (2015).

24. Bolkan, S. S. et al. Thalamic projections sustain prefrontal activity during working memory maintenance. Nat. Neurosci. 20, 987–996 (2017).

25. Rikhye, R. V, Gilra, A. & Halassa, M. M. Thalamic regulation of switching between cortical representations enables cognitive flexibility. Nat. Neurosci. 21, 1753–1763 (2018).

26. Hauer, B. E., Pagliardini, S. & Dickson, C. T. The Reuniens Nucleus of the Thalamus Has an Essential Role in Coordinating Slow-Wave Activity between Neocortex and Hippocampus. eneuro 6, ENEURO.0365-19.2019 (2019).

27. Schmitt, L. I. et al. Thalamic amplification of cortical connectivity sustains attentional control. Nature 545, 219–223 (2017).

28. Raghavachari, S. et al. Gating of human theta oscillations by a working memory task. J. Neurosci. 21, 3175–83 (2001).

29. Roberts, B. M., Hsieh, L.-T. & Ranganath, C. Oscillatory activity during maintenance of spatial and temporal information in working memory. Neuropsychologia 51, 349–57 (2013).

30. Motes, M. A. & Rypma, B. Working memory component processes: isolating BOLD signal changes. Neuroimage 49, 1933–41 (2010).

31. Riggall, A. C. & Postle, B. R. The relationship between working memory storage and elevated activity as measured with functional magnetic resonance imaging. J. Neurosci. 32, 12990–8 (2012).

32. Rottschy, C. et al. Modelling neural correlates of working memory: a coordinate-based meta-analysis. Neuroimage 60, 830–46 (2012).

33. Sreenivasan, K. K., Curtis, C. E. & D’Esposito, M. Revisiting the role of persistent neural activity during working memory. Trends Cogn. Sci. 18, 82–9 (2014).

34. Lundqvist, M., Herman, P. & Miller, E. K. Working Memory: Delay Activity, Yes! Persistent Activity? Maybe Not. J. Neurosci. 38, 7013–7019 (2018).

35. Cavanagh, S. E., Towers, J. P., Wallis, J. D., Hunt, L. T. & Kennerley, S. W. Reconciling persistent and dynamic hypotheses of working memory coding in prefrontal cortex. Nat. Commun. 9, 1–16 (2018).

36. Harvey, C. D., Coen, P. & Tank, D. W. Choice-specific sequences in parietal cortex during a virtual-navigation decision task. Nature 484, 62–68 (2012).

37. Rajan, K., Harvey, C. D. & Tank, D. W. Recurrent Network Models of Sequence Generation and Memory. Neuron 90, 128–42 (2016).

38. Park, J. C., Bae, J. W., Kim, J. & Jung, M. W. Dynamically changing neuronal activity supporting working memory for predictable and unpredictable durations. Sci. Rep. 9, 15512 (2019).

39. Murray, J. D. et al. Stable population coding for working memory coexists with heterogeneous neural dynamics in prefrontal cortex. Proc. Natl. Acad. Sci. 114, 394–399 (2017).

40. Loewenstein, Y. & Sompolinsky, H. Temporal integration by calcium dynamics in a model neuron. Nat. Neurosci. 6, 961–967 (2003).

41. Hasselmo, M. E. & Stern, C. E. Mechanisms underlying working memory for novel information. Trends Cogn. Sci. 10, 487–493 (2006).

42. Mongillo, G., Barak, O. & Tsodyks, M. SynaptiC Theory of Working Memory. Science (80-.). 319, 1543–1546 (2008).

43. Jaeger, H. Short term memory in echo state networks. GMD Rep. 152 60 (2002).

44. Vogels, T. P. & Abbott, L. F. Signal propagation and logic gating in networks of integrate-and-fire neurons. J. Neurosci. 25, 10786–95 (2005).

45. Druckmann, S. & Chklovskii, D. B. Neuronal circuits underlying persistent representations despite time varying activity. Curr. Biol. 22, 2095–103 (2012).

46. Stokes, M. G. ‘Activity-silent’ working memory in prefrontal cortex: A dynamic coding framework. Trends Cogn. Sci. 19, 394–405 (2015).

47. Baeg, E. H. et al. Dynamics of population code for working memory in the prefrontal cortex. Neuron 40, 177–188 (2003).

48. Balaguer-Ballester, E., Lapish, C. C., Seamans, J. K. & Durstewitz, D. Attracting Dynamics of Frontal Cortex Ensembles during Memory-Guided Decision-Making. PLoS Comput. Biol. 7, e1002057 (2011).

49. Leavitt, M. L., Pieper, F., Sachs, A. J. & Martinez-Trujillo, J. C. Correlated variability modifies working memory fidelity in primate prefrontal neuronal ensembles. Proc. Natl. Acad. Sci. U. S. A. 114, E2494–E2503 (2017).

50. Rigotti, M. et al. The importance of mixed selectivity in complex cognitive tasks. Nature 497, 585–590 (2013).

51. Parthasarathy, A. et al. Mixed selectivity morphs population codes in prefrontal cortex. Nat. Neurosci. 20, 1770–1779 (2017).

52. Balaguer-Ballester, E., Nogueira, R., Abofalia, J. M., Moreno-Bote, R. & Sanchez-Vives, M. V. Representation of foreseeable choice outcomes in orbitofrontal cortex triplet-wise interactions. PLOS Comput. Biol. 16, e1007862 (2020).

53. Spaak, E., Watanabe, K., Funahashi, S. & Stokes, M. G. Stable and Dynamic Coding for Working Memory in Primate Prefrontal Cortex. J. Neurosci. 37, 6503–6516 (2017).

54. Watanabe, T. & Niki, H. Hippocampal unit activity and delayed response in the monkey. Brain Res. 325, 241–254 (1985).

55. Navawongse, R. & Eichenbaum, H. Distinct pathways for rule-based retrieval and spatial mapping of memory representations in hippocampal neurons. J. Neurosci. 33, 1002–1013 (2013).

56. Jones, M. W. & Wilson, M. a. Theta rhythms coordinate hippocampal-prefrontal interactions in a spatial memory task. PLoS Biol. 3, e402 (2005).

57. Benchenane, K. et al. Coherent theta oscillations and reorganization of spike timing in the hippocampal-prefrontal network upon learning. Neuron 66, 921–36 (2010).

58. O’Neill, P. K., Gordon, J. A. & Sigurdsson, T. Theta oscillations in the medial prefrontal cortex are modulated by spatial working memory and synchronize with the hippocampus through its ventral subregion. J. Neurosci. 33, 14211–14224 (2013).

59. Myroshnychenko, M., Seamans, J. K., Phillips, A. G. & Lapish, C. C. Temporal Dynamics of Hippocampal and Medial Prefrontal Cortex Interactions during the Delay Period of a Working Memory-Guided Foraging Task. Cereb. Cortex 27, 5331–5342 (2017).

60. Place, R., Farovik, A., Brockmann, M. & Eichenbaum, H. Bidirectional prefrontal-hippocampal interactions support context-guided memory. Nat. Neurosci. 19, 992–994 (2016).

61. Brincat, S. L. & Miller, E. K. Frequency-specific hippocampal-prefrontal interactions during associative learning. Nat. Neurosci. 18, 576–581 (2015).

62. Liu, Y., Brincat, S. L., Miller, E. K. & Hasselmo, M. E. A Geometric Characterization of Population Coding in the Prefrontal Cortex and Hippocampus during a Paired-Associate Learning Task. J. Cogn. Neurosci. 32, 1455–1465 (2020).

63. Meyers, E. M., Freedman, D. J., Kreiman, G., Miller, E. K. & Poggio, T. Dynamic population coding of category information in inferior temporal and prefrontal cortex. J. Neurophysiol. 100, 1407–1419 (2008).

64. Spellman, T. et al. Hippocampal-prefrontal input supports spatial encoding in working memory. Nature 522, 309–314 (2015).

65. Maharjan, D. M., Dai, Y. Y., Glantz, E. H. & Jadhav, S. P. Disruption of dorsal hippocampal – prefrontal interactions using chemogenetic inactivation impairs spatial learning. Neurobiol. Learn. Mem. 155, 351–360 (2018).

66. Schmidt, X. B., Duin, A. A. & Redish, A. D. Disrupting the medial prefrontal cortex alters hippocampal sequences during deliberative decision making. J. Neurophysiol. 121, 1981–2000 (2019).

67. Krzanowski, W. J. Principles of Multivariate Analysis: A User’s Perspective. (Oxford University Press, 2000).

68. Durstewitz, D. Advanced Data Analysis in Neuroscience. (Springer, 2017).

69. Yu, B. M. et al. Gaussian-process factor analysis for low-dimensional single-trial analysis of neural population activity. J. Neurophysiol. 102, 614–635 (2009).

70. Xia, M., Liu, T., Bai, W., Zheng, X. & Tian, X. Information transmission in HPC-PFC network for spatial working memory in rat. Behav. Brain Res. 356, 170–178 (2019).

71. Liu, T., Bai, W., Xia, M. & Tian, X. Directional hippocampal-prefrontal interactions during working memory. Behav. Brain Res. 338, 1–8 (2018).

72. Kucewicz, M. T., Tricklebank, M. D., Bogacz, R. & Jones, M. W. Dysfunctional prefrontal cortical network activity and interactions following cannabinoid receptor activation. J. Neurosci. 31, 15560–15568 (2011).

73. Izaki, Y., Takita, M. & Akema, T. Specific role of the posterior dorsal hippocampus-prefrontal cortex in short-term working memory. Eur. J. Neurosci. 27, 3029–3034 (2008).

74. Yoon, T., Okada, J., Jung, M. W. & Kim, J. J. Prefrontal cortex and hippocampus subserve different components of working memory in rats. Learn. Mem. 15, 97–105 (2008).

75. Churchwell, J. C., Morris, A. M., Musso, N. D. & Kesner, R. P. Prefrontal and hippocampal contributions to encoding and retrieval of spatial memory. Neurobiol. Learn. Mem. 93, 415–421 (2010).

76. Newmark, R. E., Schon, K., Ross, R. S. & Stern, C. E. Contributions of the Hippocampal Subfields and Entorhinal Cortex to Disambiguation During Working Memory. Hippocampus 23, 467–475 (2013).

77. Gluth, S., Sommer, T., Rieskamp, J. & Büchel, C. Effective Connectivity between Hippocampus and Ventromedial Prefrontal Cortex Controls Preferential Choices from Memory. Neuron 86, 1078–1090 (2015).

78. Roberts, B. M., Libby, L. A., Inhoff, M. C. & Ranganath, C. Brain activity related to working memory for temporal order and object information. Behav. Brain Res. 354, 55–63 (2018).

79. Gunseli, E. & Aly, M. Preparation for upcoming attentional states in the hippocampus and medial prefrontal cortex. Elife 9, 1–33 (2020).

80. Laroche, S., Davis, S. & Jay, T. M. Plasticity at hippocampal to prefrontal cortex synapses: Dual roles in working memory and consolidation. Hippocampus 10, 438–446 (2000).

81. Thierry, A. M., Gioanni, Y., Dégénétais, E. & Glowinski, J. Hippocampo-prefrontal cortex pathway: Anatomical and electrophysiological characteristics. Hippocampus 10, 411–419 (2000).

82. Hoover, W. B. & Vertes, R. P. Anatomical analysis of afferent projections to the medial prefrontal cortex in the rat. Brain Struct. Funct. 212, 149–179 (2007).

83. Gilmartin, M. R., Miyawaki, H., Helmstetter, F. J. & Diba, K. Prefrontal Activity Links Nonoverlapping Events in Memory. J. Neurosci. 33, 10910–10914 (2013).

84. Canetta, S. et al. Differential Synaptic Dynamics and Circuit Connectivity of Hippocampal and Thalamic Inputs to the Prefrontal Cortex. Cereb. Cortex Commun. 1, (2020).

85. Holtmaat, A. & Caroni, P. Functional and structural underpinnings of neuronal assembly formation in learning. Nat. Neurosci. 19, 1–10 (2016).

86. Fujisawa, S. & Buzsáki, G. A 4 Hz Oscillation Adaptively Synchronizes Prefrontal, VTA, and Hippocampal Activities. Neuron 72, 153–165 (2011).

87. Floresco, S. B., Seamans, J. K. & Phillips, A. G. Selective Roles for Hippocampal, Prefrontal Cortical, and Ventral Striatal Circuits in Radial-Arm Maze Tasks With or Without a Delay. J. Neurosci. 17, 1880–1890 (1997).

88. Meyer-Lindenberg, A. S. et al. Regionally specific disturbance of dorsolateral prefrontal-hippocampal functional connectivity in schizophrenia. Arch. Gen. Psychiatry 62, 379–386 (2005).

89. Sigurdsson, T., Stark, K. L., Karayiorgou, M., Gogos, J. a & Gordon, J. a. Impaired hippocampal-prefrontal synchrony in a genetic mouse model of schizophrenia. Nature 464, 763–7 (2010).

90. Sales-Carbonell, C. et al. Striatal GABAergic and cortical glutamatergic neurons mediate contrasting effects of cannabinoids on cortical network synchrony. Proc. Natl. Acad. Sci. U. S. A. 110, (2013).

91. Sandler, R. A., Fetterhoff, D., Hampson, R. E., Deadwyler, S. A. & Marmarelis, V. Z. Cannabinoids disrupt memory encoding by functionally isolating hippocampal CA1 from CA3. PLOS Comput. Biol. 13, e1005624 (2017).

92. Paxinos, G. & Watson, C. The Rat Brain in Stereotaxic Coordinates, Sixth Edition: Hard Cover Edition. (Academic Press, 2007).

93. Taylor, C. C. Bootstrap Choice of the Smoothing Parameter in Kernel Density Estimation. Biometrika 76, 705–712 (1989).

94. Shimazaki, H. & Shinomoto, S. Kernel bandwidth optimization in spike rate estimation. J. Comput. Neurosci. 29, 171–182 (2010).

95. Hastie, T., Tibshirani, R. & Friedman, J. The Elements of Statistical Learning: Data Mining, Inference, and Prediction. (Springer, 2011).

96. Stokes, M. G. et al. Dynamic coding for cognitive control in prefrontal cortex. Neuron 78, 364–375 (2013).

97. Cunningham, J. P. & Yu, B. M. Dimensionality reduction for large-scale neural recordings. Nat. Neurosci. 17, 1500–1509 (2014).

98. Lopes-dos-Santos, V., Ribeiro, S. & Tort, A. B. L. Detecting cell assemblies in large neuronal populations. J. Neurosci. Methods 220, 149–166 (2013).

